# Patient-derived Induced Pluripotent Stem Cells as a Model to Study Frontotemporal Dementia Pathologies

**DOI:** 10.1101/2024.12.11.628042

**Authors:** Sonia Infante-Tadeo, Diane L. Barber

## Abstract

The neurodegenerative disorder Frontotemporal Dementia (FTD) can be caused by a repeat expansion (GGGGCC; G4C2) in C9orf72. The function of wild-type C9orf72 and the mechanism by which the C9orf72-G4C2 mutation causes FTD, however, remain unresolved. Diverse disease models including human brain samples and differentiated neurons from patient-derived induced pluripotent stem cells (iPSCs) identified some hallmarks associated with FTD, but these models have limitations, including biopsies capturing only a static snapshot of dynamic processes and differentiated neurons being labor-intensive, costly, and post-mitotic. We find that patient-derived iPSCs, without being differentiated into neurons, exhibit established FTD hallmarks, including increased lysosome pH, decreased lysosomal cathepsin activity, cytosolic TDP-43 proteinopathy, and increased nuclear TFEB. Moreover, lowering lysosome pH in FTD iPSCs mitigates TDP-43 proteinopathy, suggesting a key role for lysosome dysfunction. RNA-seq reveals dysregulated transcripts in FTD iPSCs affecting calcium signaling, cell death, synaptic function, and neuronal development. We confirm differences in protein expression for some dysregulated genes not previously linked to FTD, including CNTFR (neuronal survival), Annexin A2 (anti-apoptotic), NANOG (neuronal development), and moesin (cytoskeletal dynamics). Our findings underscore the potential of FTD iPSCs as a model for studying FTD cellular pathology and for drug screening to identify therapeutics.

**SIGNIFICANCE STATEMENT:**

- Understanding the cellular pathology of Frontotemporal Dementia linked to a GGGGCC expansion in the C9orf72 gene remains a challenge.
- This study shows that undifferentiated patient-derived iPSCs exhibit hallmark FTD characteristics, including lysosome dysfunction and TDP-43 proteinopathy, and identifies dysregulated genes related to neurodegeneration.
- These findings highlight patient-derived iPSCs as a valuable model for studying FTD pathology and for drug screening, potentially guiding future research in therapeutic development.

## INTRODUCTION

Mutations in C9orf72 are the most common inherited genetic cause of Frontotemporal Dementia (FTD) as well as Amyotrophic Lateral Sclerosis (ALS) (Sonobe et al., 2023; Todd et al., 2023). FTD, a dementia related to Alzheimer’s disease (AD), commonly affects individuals under 65 years old. Genetic factors contribute to 30–40% of FTD cases, with mutations in C9orf72 being the most prevalent, but FTD is also associated with mutations in progranulin (GRN) or tau (MAPT) (Greaves and Rohrer, 2019). Mutations in C9orf72 associated with FTD are typically a hexanucleotide (GGGGCC [G4C2]) repeat expansion (DeJesus-Hernandez et al., 2011; Renton et al., 2011). While healthy individuals typically possess fewer than 20 G4C2 repeats, expansions from 30 to more than 100 repeats are linked to ALS and FTD (Rademakers et al., 2012; Babic Leko et al., 2019). Common cellular abnormalities associated with C9orf72-G4C2 include a disrupted endolysosomal pathway, increased lysosomal pH (pHlys), reduced lysosomal hydrolyze activity, and TDP-43 proteinopathy, which collectively contribute to cell damage (Ling et al., 2013; Amick and Ferguson, 2017). Although the role of wild-type C9orf72 in maintaining lysosome balance is acknowledged (Amick and Ferguson, 2017), the mechanisms whereby mutant C9orf72-G4C2 leads to impaired lysosome biology, toxicity, and neurodegeneration remain unclear (Babic Leko et al., 2019).

Our current understanding of FTD pathogenesis comes from different experimental models, each with advantages and limitations. Post-mortem brain samples from FTD patients provide *in situ* analysis and tissue level pathologies but capture only a static snapshot of dynamic processes and are not amenable to genetic manipulation for mechanistic analysis or drug screening to identify potential therapeutics (Halliday et al., 2011; Irwin et al., 2015). Models in mice (Chew et al., 2015; Czuppa et al., 2022; Lopez-Herdoiza et al., 2023) and other organisms including zebrafish (Shaw et al., 2018) and Drosophila (Mizielinska et al., 2014) provide systems-level and genetic manipulation but with species differences that do not effectively recapitulate human FTD (Neumann and Mackenzie, 2019; Ahmed et al., 2023; White et al., 2024). An increasingly common cellular model is neurons differentiated from induced pluripotent stem cells (iPSCs) generated from patients with FTD (Lee and Huang, 2017; Beckers and Van Damme, 2024). Using differentiated neurons revealed several key findings, including changes in lysosomal cathepsin activity (Lee and Huang, 2017; Valdez et al., 2017), aberrant cytosolic aggregates of TDP-43, and dysregulated TFEB and TFE3, which are transcription factors regulating lysosome biogenesis. However, despite recently improved protocols to differentiate neurons from iPSCs (Espuny-Camacho et al., 2013; Raitano et al., 2015; Qi et al., 2017), the procedure remains time-extensive, costly, and the most streamlined methods require genetic manipulations (Lara Flores et al., 2023). Moreover, as post-mitotic cells, terminally differentiated neurons cannot be propagated for continual use. To our knowledge, the use of FTD patient-derived iPSCs as a tractable experimental model without differentiation to neurons has received limited attention, indicating a potential avenue for further investigation (Selvaraj et al., 2017; Lines et al., 2020). In contrast, ALS patient-derived iPSCs have been used as a model system for examining epigenetic perturbations caused by C9orf72 mutations (Esanov et al., 2016).

In this report we tested the feasibility of using iPSCs as a tractable model for studying FTD cellular pathologies. We compared iPSCs derived from a patient with FTD carrying a C9orf72-G4C2 expansion with control wild-type iPSCs from a gender, age, and race matched non-affected individual. Defective lysosomal acidification causes neuroinflammation and neurodegeneration through a variety of detrimental cellular pathways (Quick et al., 2023). We found that FTD iPSCs have hallmarks of FTD-associated pathologies, including increased pHlys, decreased cathepsin activity, TDP-43 proteinopathy with cytoplasmic aggregates, and nuclear accumulation of TFEB, which are consistent with findings in patient tissues (Smeele et al., 2024) and neurons differentiated from iPSCs (Lee and Huang, 2017; Valdez et al., 2017). Furthermore, RNA-seq analysis identified transcriptomics differences in FTD iPSCs compared with WT iPSCs, including dysregulated genes involved in neuroinflammation, calcium signaling, synaptic function, and neuronal development not previously linked to FTD. The significance of our findings includes 1) confirming that dysregulated cell functions with C9orf72-G4C2 are not limited to neurons, 2) identifying new regulators of dysregulated cell functions with FTD, and 3) showing the promise of using FTD iPSCs as a previously unreported experimental model for investigating FTD cellular pathologies and as a platform for drug screening to identify therapeutics for treating FTD (Vo et al., 2024).

## RESULTS

Our study used iPSCs sourced from the National Institute of Neurological Disorders and Stroke (NINDS), including cells derived from an FTD-affected 48-year-old Caucasian male with a C9orf72-G4C2 expansion (FTD iPSCs) and a non-affected age, gender, and race matched individual (wild-type [WT] iPSCs). We confirmed the pluripotent state of both cell lines with standard markers and found no differences between WT compared with FTD iPSCs for the characteristics we determined, including colony morphology (Supplemental Figure S1A), E-cadherin abundance and localization at cell-cell junctions (Supplemental Figure S1B, E), and predominantly nuclear localization of stem cell pluripotency transcription factors OCT4 and SOX2 (Supplemental Figure S1C, D) as well as total abundance of OCT4 (Supplemental Figure S1E).

### FTD iPSCs have increased lysosome pH

Although the exact mechanisms triggering FTD pathologies with C9orf72-G4C2 remain unclear, an aberrantly high pHlys is reported with early-stages of different types of neurodegeneration (Wolfe et al., 2013; Nixon, 2020; Chin et al., 2022). We therefore quantified pHlys by generating WT and FTD iPSCs stably expressing a genetically encoded and ratiometric fluorescent biosensor, pHLARE, which we previously described accurately measures pHlys within the range of pH 4.0 to 6.5 (Webb et al., 2021). pHLARE encodes the Lysosomal Associated Membrane Protein 1 (LAMP1) tagged at the lumenal domain with superfolder GFP and at the cytosolic domain with mCherry. We stably expressed in iPSC lines pHLARE containing an EF-1α promoter inserted into the AAVS1 harbor locus, which is a site commonly used for stable expression in stem cells (Smith et al., 2008), by using MaxCyte ATX electroporation and FACS sorting for mCherry. Measuring changes in ratiometric pHLARE calibrated to pH values with nigericin buffers at the end of each determination as described (Webb et al., 2021), we found that the pHlys of FTD iPSCs, 5.0 ± 0.2, was significantly higher compared with WT iPSCs, 4.4 ± 0.2 (Figure 1A, B). To validate our findings obtained with pHLARE, we performed ratiometric fluorescence imaging using the pH-sensitive dye Oregon Green 488 Dextran. Consistent with the pHLARE measurements, WT iPSCs showed an average pH of 5.3 ± 0.8, whereas FTD iPSCs showed an increased average pH of around 6.1 ± 0.8 (Figure S2A). The slightly higher pH detected with Oregon Green 488 Dextran compared with pHLARE likely reflects its lower specificity for lysosomes, as the dye is also distributed in late endosomes (Halcrow et al., 2021). Compared with previous findings, a higher pHlys is also seen in ALS iPSC-derived neurons (Lo and Zeng, 2023), in fibroblasts from Parkinson’s Disease (PD) patients (Colacurcio and Nixon, 2016), and in iPSCs-derived neurons (i-neurons) from a patient carrying a common FTD-linked mutation in the GRN locus (Elia et al., 2023).

**FIGURE 1:**
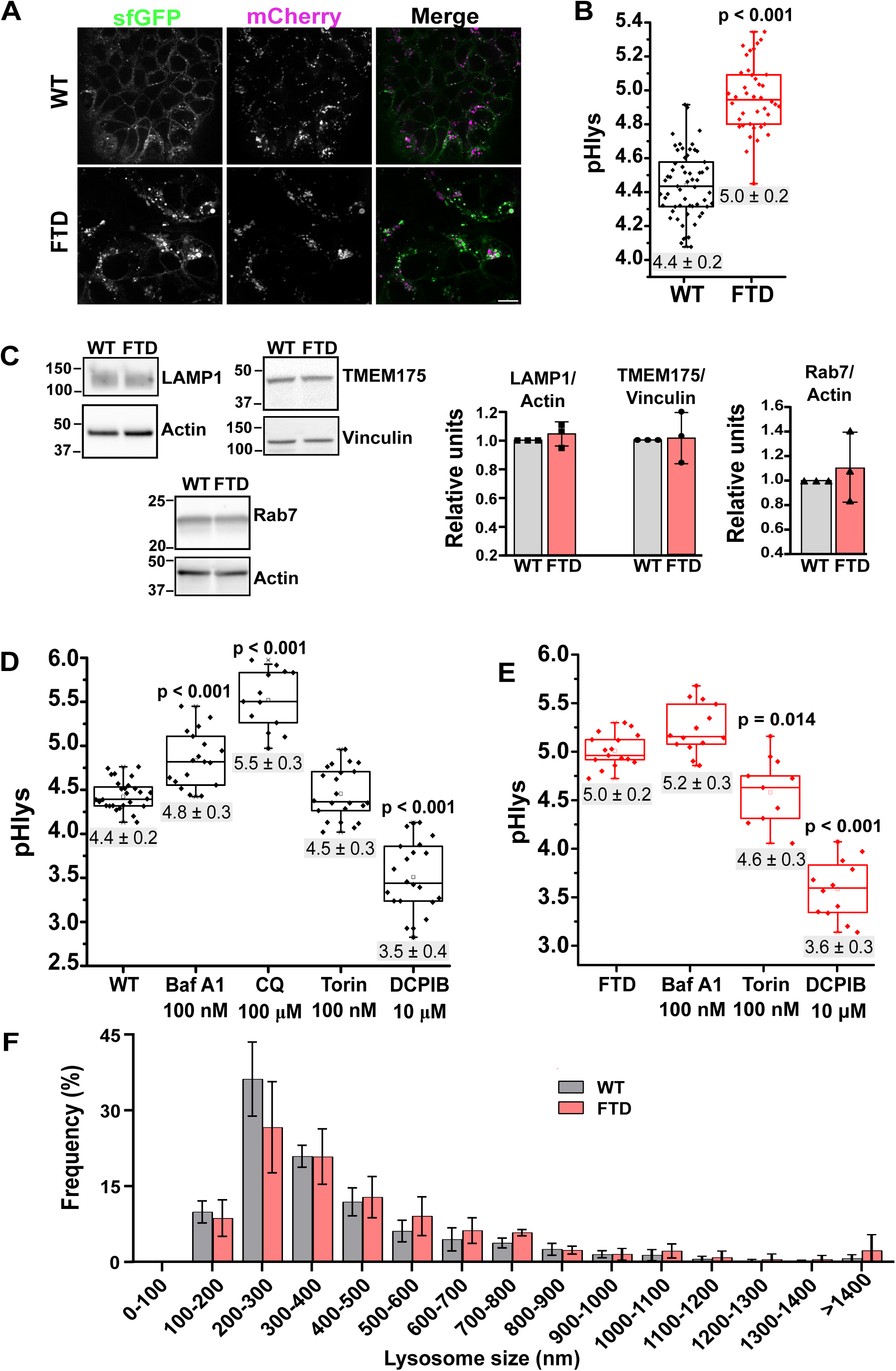
Lysosome Characteristics in FTD iPSCs (A) Confocal images of WT and FTD iPSCs expressing pHLARE, showing sfGFP, mCherry, and merged fluorescence signals. Scale bar 15 μm. (B) Quantified pHlys in WT and FTD iPSCs obtained from pHLARE sfGFP/mCherry ratios calibrated with nigericin-containing buffers of known pH values. Box plots show median, first, and third quartile, with whiskers extending to observations within 1.5 times the interquartile range, with data points indicating individual cells (WT 60 cells; FTD 45 cells) obtained from 5 separate cell preparations. Statistical analysis by Dunnett’s multiple comparison test. (C) Representative immunoblots and quantified data from 3 separate cell preparations for abundance of TMEM175, LAMP1 and Rab7 in lysates of WT and FTD iPSC normalized to loading controls of either vinculin or actin (D, E) The pHlys of WT (D) and FTD (E) iPSCs after treatment with different chemical reagents. Data were expressed as described in B and obtained from three separate cell preparations. (F) Lysosome size determined by measuring ca. 1000 lysosomes per cell line in four cell preparations.

Despite the higher pHlys we observed in FTD iPSCs, immunoblotting total cell lysates revealed that WT and FTD iPSC had a similar abundance of proteins reported to regulate pHlys, including LAMP1 and Transmembrane protein 175 (TMEM175) (Figure 1C). TMEM175 was recently identified as a lysosomal K^+^-H^+^ channel with a pivotal role in maintaining lysosomal membrane potential and pHlys stability (Hu et al., 2022; Tang et al., 2023). LAMP1 was subsequently shown to bind to TMEM175 and regulate channel activity and pHlys (Zhang et al., 2023). Additionally, despite defective endolysosomal trafficking reported as an FTD pathology (Farg et al., 2014) (Shi et al., 2018; Shao et al., 2022), WT and FTD iPSCs had a similar abundance of Ras-related protein Rab7, which confers maturation of late endosomes into lysosomes and lysosomal trafficking. These data are consistent with previous findings indicating no significant differences in LAMP1 expression within the frontal grey matter of FTD-C9orf72 individuals compared with control samples (Marian et al., 2023) or progranulin-deficient FTD cells (Valdez et al., 2017), as well as no significant difference in Rab7 expression in C9orf72 knockout mice (Todd et al., 2023).

We could experimentally change pHlys in both WT and FTD iPSCs. The pHlys of WT iPSCs increased with Bafilomycin A1 (Baf A1; 100 nM), an inhibitor of the vacuolar V-ATPase, the main proton pump in lysosomes (Tu et al., 2008; Casey et al., 2010), and chloroquine (CQ; 100 μM), an alkalinizing lysosomotropic compound (Levine and Kroemer, 2008; Tu et al., 2008), but did not change with Torin-1 (100 nM), an ATP-competitive inhibitor of mammalian target of rapamycin complex 1 (mTORC1) activity that lowers pHlys in somatic cells (Thoreen et al., 2009; Webb et al., 2021) (Figure 1D). In contrast, in FTD iPSCs Torin-1 lowered pHlys but Bafilomycin had no effect (Figure 1E). We also found that the pHlys of both WT and FTD pHlys was decreased when cells were treated with 4-(2-(6,7-Dichloro-2-cyclopentylindan-1-on-5-yl)ethyl)benzene-1,3-diol (DCPIB; 10 μM), which is reported to be a selective inhibitor of volume-regulated anion channels (VRACs) or swelling-activated chloride channels (Decher et al., 2001; Sardini et al., 2003) (Figure 1D and 1E). In contrast to DCPIB inhibiting VRACs and chloride channels, Hu and colleagues (Hu et al., 2022) reported that DCPIB targets TMEM175 to increase channel activity and pHlys in HEK293T cells.

Although FTD iPSCs had a higher pHlys compared with WT, their cytosolic pH (pHi) was not different (WT 7.57 ± 0.04, FTD 7.55 ± 0.04) (Supplemental Figure S2B), determined in cell populations loaded with the fluorescence pH sensitive dye 2′,7′-Bis(2-carboxyethyl)-5(6)-carboxyfluorescein (BCECF). Additionally, DCPIB had no effect on pHi of either WT or FTD iPSCs (Supplemental Figure S2B), suggesting a selective effect on pHlys. These findings are in contrast with studies, albeit limited, indicating that decreased pHi was associated with AD and other neurodegenerative diseases (Fang et al., 2010; Majdi et al., 2016), although to our knowledge differences in pHi with FTD have not been reported.

Lysosome size is noted to be variable but generally ranges between 150—2000 nm, as determined in epithelial cells (Bandyopadhyay et al., 2014; Webb et al., 2021; Siddiqui and Bhatt, 2023). We found that WT and FTD had a mostly similar profile of lysosome size, including a mean size of 400 ± 235 nm for WT iPSCs and 500 ± 410 nm for FTD iPSCs. However, WT iPSCs had more smaller lysosomes between 200—300 nm and FTD iPSCs had more larger lysosomes >1400 nm (Figure 1F). A similar finding of increased proportion of enlarged lysosomes is reported using transmission electron microscopy of C9orf72 iPSC-derived motor neurons (Beckers et al., 2023). Together, our findings indicate that compared with WT iPSCs, FTD iPSCs had a higher pHlys and somewhat larger lysosome size but no difference in pHi or the expression of LAMP1, TMEM175, and Rab7.

### Proteolytic activity and Cathepsin B activity and expression are decreased in FTD iPSCs

Lysosomal degradation relies on over 50 hydrolases, including many cathepsins with optimal activity in an acidic environment (Wolfe et al., 2013). We evaluated lysosomal hydrolase activity by staining cells with DQ™-Red BSA, a fluorogenic substrate that is internalized through endocytosis and transported to endosomes and lysosomes. In acidic compartments the BSA is degraded by lysosomal hydrolases, releasing BODIPY™ dye molecules. To confirm this behavior of DQ™-Red BSA, we showed significantly decreased fluorescence in WT iPSCs treated with Bafilomycin A1 (100 nM, 18 h) (Figure 2A), which increased pHlys (Figure 1D). In control FTD cells we found decreased fluorescence compared with control WT iPSCs (Figure 2A), indicating a lower proteolytic degradation. However, fluorescence increased to that seen in control WT iPSCs when we treated FTD iPSCs with DCPIB or Torin (Figure 2A), which lowered pHlys (Figure 1E).

**FIGURE 2:**
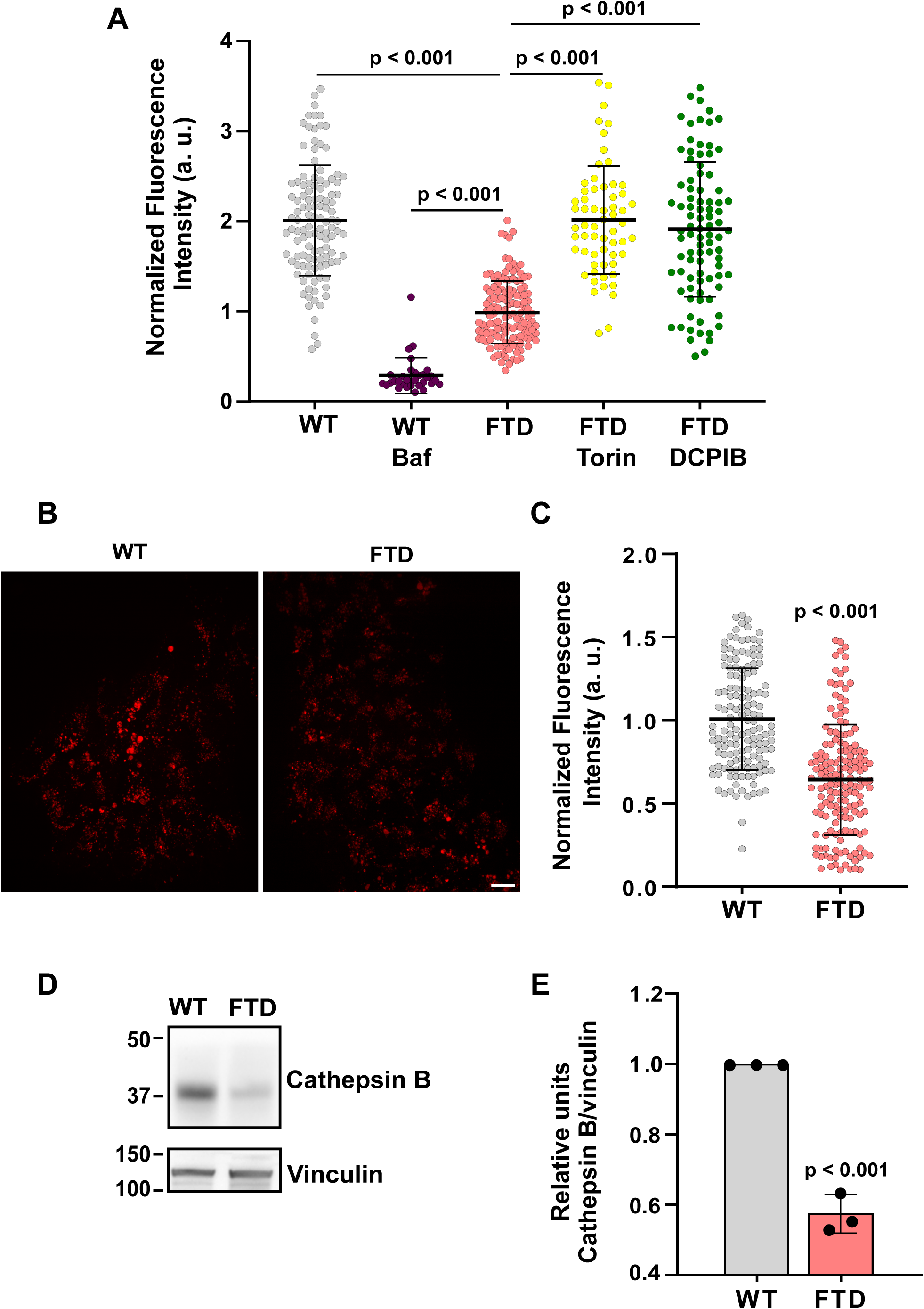
Hydrolases and Cathepsin B Activity and Expression. (A) DQ™-Red BSA assay of iPSCs upon treatment with different chemical drugs that modify the pHlys. DQ-BSA intensity was calculated by collecting at least 30 different ROIs for each biological replicate. (B) Representative fluorescence microscopy images of cells stained with Magic Red, an indicator of Cathepsin B activity. Scale bar 30 μm. (C) Quantified fluorescence intensity (a. u.) of Magic Red staining. Each dot represents the fluorescence intensity measured in a randomly selected ROI, which may cover one or more cells. (D) Immunoblot of total cell lysates probed with antibodies for Cathepsin B, and for vinculin as a loading control. (E) Quantification of Cathepsin B immunoblotting (mean ± SD) determined from 3 separate cell preparations. Data were presented as normalized values and error bars denote standard deviation from N=3 biological replicates, relative to FTD control. Statistical significance was analyzed using a one-way ANOVA and P values were shown in the figure.

Consistent with the higher pHlys in FTD compared with WT iPSCs and the reduced hydrolase activity observed in FTD iPSCs, the latter also had decreased cathepsin B activity (Figure 2B, C), determined by staining with Magic Red, a fluorogenic reporter specifically cleaved by cathepsin B in lysosomes. FTD iPSCs, in addition to having decreased cathepsin B activity, also had decreased cathepsin B abundance compared with WT, determined by immunoblotting total cell lysates (Figure 2D, E). Although decreased activity of lysosomal cathepsin D is seen in iPSC-derived cortical neurons from FTD patients harboring progranulin mutations (Valdez et al., 2017), to our knowledge a link between C9orf72-G4C2 and decreased cathepsin B activity has not been reported. However, decreased cathepsin B abundance is seen in the cerebrospinal fluid of patients with primary progressive aphasia, a language variant of FTD (Swift et al., 2024). Additionally, inhibiting cathepsin B activity or its loss is reported to promote development of AD and neurodegeneration (Kos et al., 2022). Additionally, previous studies highlighted that bone marrow-derived cells exposed to pHlys-elevating chemicals have a notable reduction in both cathepsin B and D protein abundance (Xu et al., 2024). Together, our data confirm that a higher pHlys correlates with decreased total hydrolase activity, determined with DQ™-Red BSA, and specifically cathepsin B activity, determined with Magic Red.

### Cytosolic protein aggregates, including TDP-43, are increased in FTD iPSCs

Aberrant cytosolic protein aggregates is a hallmark of neurodegenerative diseases such as AD, PD, ALS, and FTD (Taylor et al., 2002; Neumann et al., 2006). We scored for global protein aggregates by using the PROTEOSTAT® Aggresome Detection Reagent, a molecular rotor dye designed to interact with and disrupt the cross-beta spine of quaternary protein structures commonly found in misfolded and aggregated proteins. In WT iPSCs the cytoplasmic fluorescence intensity remained relatively low with a mean of 250 arbitrary units (a.u.), but increased with Bafilomycin A treatment (100 nM, 2 h) to a mean of 400 a.u. (Figure 3A–B). In FTD iPSCs the mean of 500 a.u. was greater than WT iPSCs, both control and treated with Bafilomycin A (Figure 3A–B).

**FIGURE 3:**
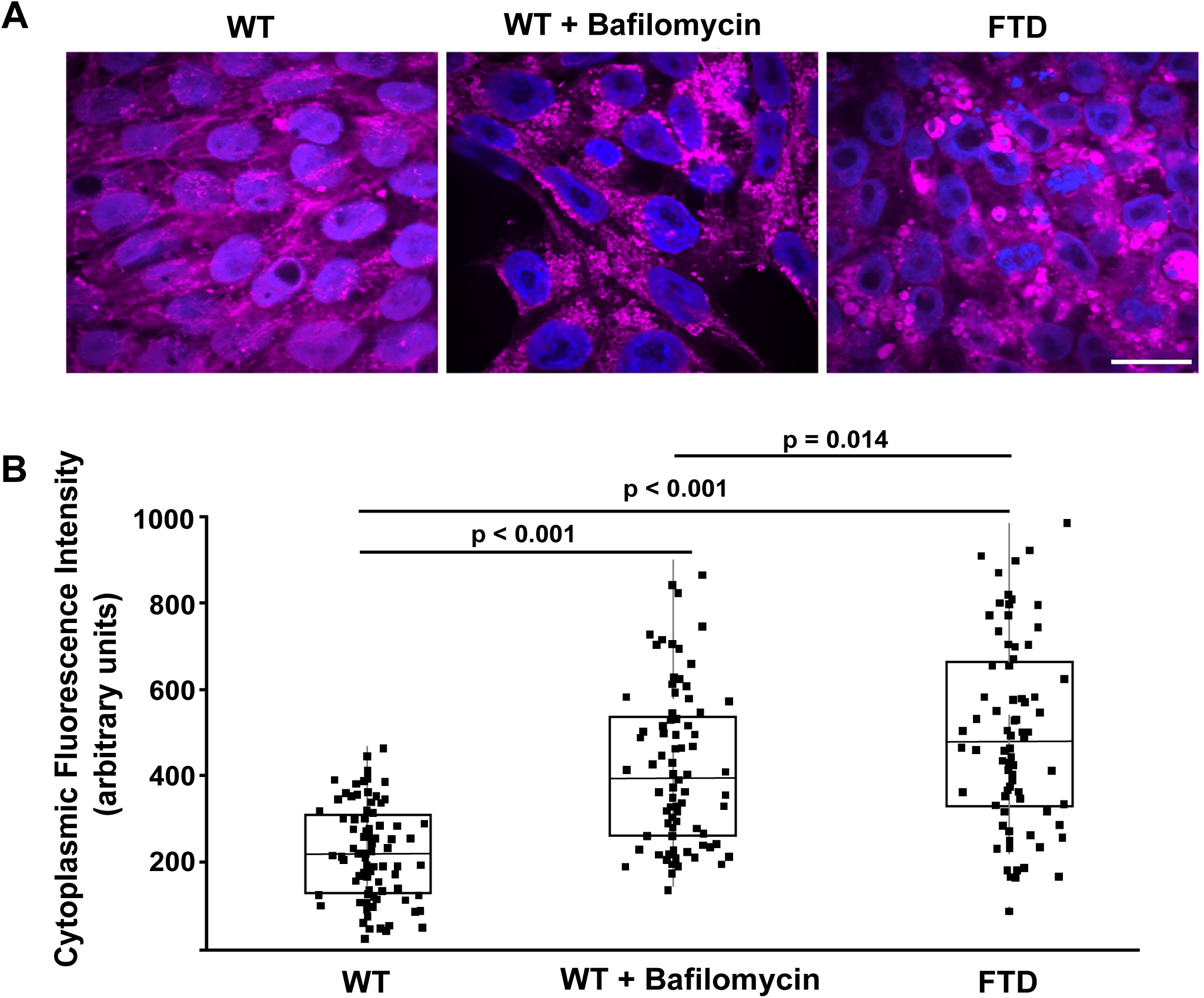
ProteoStat Labeling and Aggregate Quantification. (A) Representative confocal images of WT iPSCs, untreated and treated with Bafilomycin, and FTD iPSCs stained with ProteoStat® (purple) and Hoechst 33342 (blue). Scale bar 10 μm. (B) Quantification of fluorescence intensities of cytosolic ProteoStat® determined from images as show in (A). Box plots show median, first, and third quartile, with whiskers extending to observations within 1.5 times the interquartile range, with data points indicating individual cells obtained from three separate cell preparations. Statistical analysis by Tukey– Kramer HSD.

To further analyze the proteinopathy in FTD iPSCs, we immunolabeled for TAR DNA-binding protein 43 (TDP-43), a multifunctional DNA/RNA-binding protein found in both the nucleus and cytoplasm (Babic Leko et al., 2019; Lines et al., 2020; Ratti et al., 2020). Decreased nuclear TDP-43 and increased cytosolic TDP-43 aggregates are a common FTD pathology seen in a mouse model harboring mutant C9orf72 (Shao et al., 2022), in affected neurons of patients with FTD (Neumann et al., 2006; Beckers and Van Damme, 2024), in post-mortem brain tissue from FTD patients with C9orf72 mutations (Marian et al., 2023), and in neurons differentiated from iPSCs generated from a patient with FTD having decreased PGRN expression (Valdez et al., 2017). Consistent with these findings, immunolabeling showed that FTD iPSCs had a lower nuclear to cytoplasmic ratio of TDP-43 and visibly more cytosolic TDP-43 aggregates (Figure 4A–C) compared with WT iPSCs; however, immunoblotting indicated that the abundance of total TDP-43 was not different in the two cell lines (Figure 4D).

**FIGURE 4:**
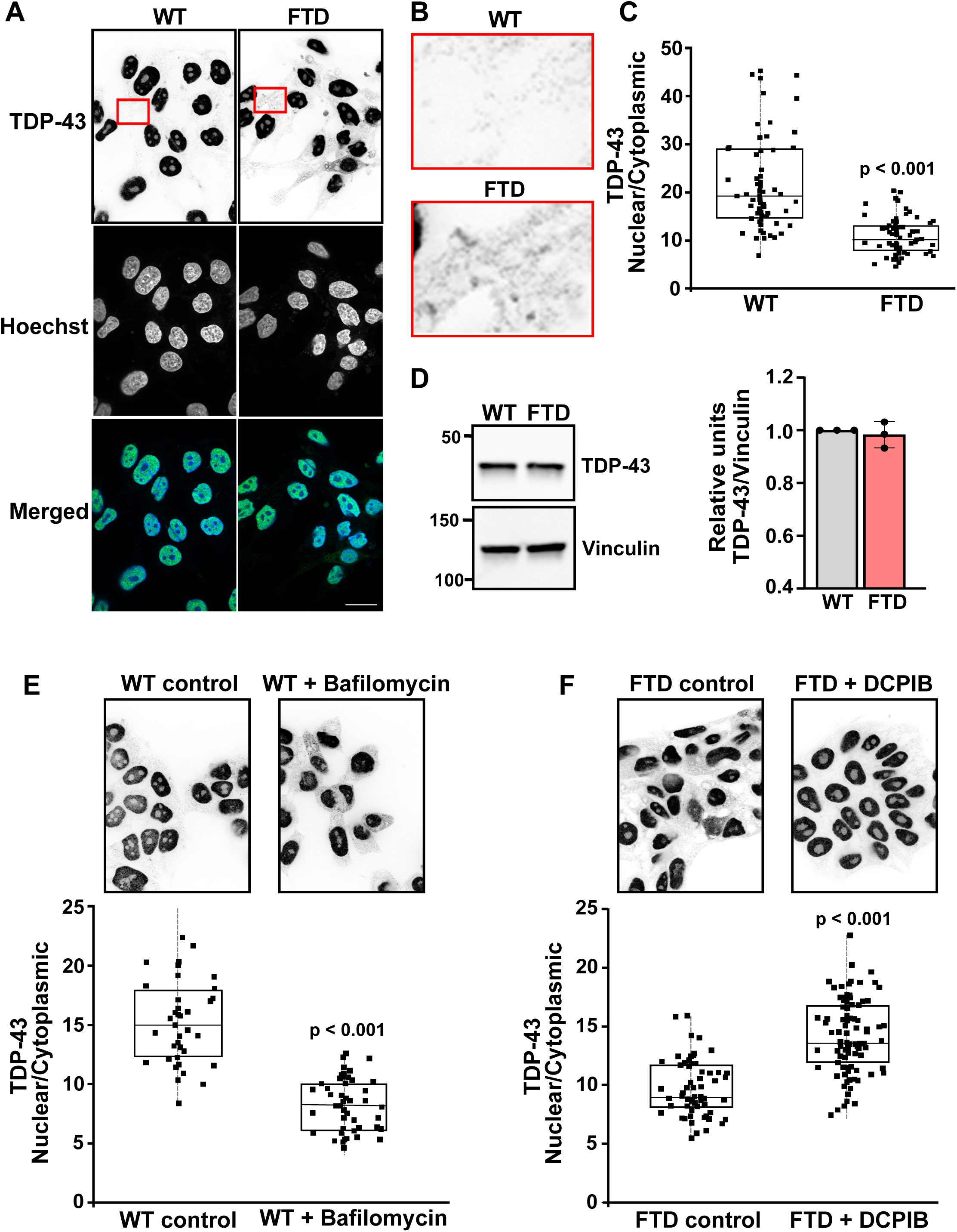
TDP-43 Localization and Expression. (A) Inverted confocal images of WT and FTD iPSCs immunolabeled with TDP-43 and stained with Hoechst 33342. Scale bar 10 μm. (B) Higher magnification of the red-highlighted areas in (A) of TDP-43 immunolabeling. (C) Quantification of the nuclear/cytoplasmic ratio of TDP-43 fluorescence in WT and FTD iPSCs determined from images as shown in (A) with each dot representing the value from a single cell. Data were presented as box plots with whiskers representing minimum to maximum values. Statistical significance was determined using an unpaired t-test (p < 0.001). (D) Representative immunoblots of TDP-43 with vinculin as a loading control and quantification of TDP-43 expression normalized to vinculin for immunoblots from three separate cell preparations. (E, F) Inverted confocal images of TDP-43 immunolabeling and quantification of nuclear/cytoplasmic ratio of TDP-43 in WT iPSCs untreated (Control) and treated with Bafilomycin (100 nM, 18h) (E) and FTD iPSCs untreated (Control) and treated with DCPIB (10 μM, 18h) (F) obtained from three separate cell preparations.

Disrupted endolysosomal trafficking is suggested to be a major determinant in TDP-43 proteinopathy (Shao et al., 2022); however a role for increased pHlys is also indicated in TDP-43 proteinopathy as seen in HeLa cells treated with Bafilomycin A (Tanaka et al., 2023). Consistent with Bafilomycin A increasing pHlys in WT iPSCs, it also decreased the nuclear to cytoplasmic ratio of TDP-43 and enhanced cytosolic TDP-43 aggregates (Figure 4E). Also, consistent with DCPIB decreasing pHlys in FTD iPSCs, it increased the nuclear to cytoplasmic ratio of TDP-43 and attenuated cytosolic TDP-43 aggregates compared with untreated control FTD cells (Figure 4F). The TDP-43 nuclear to cytoplasmic ratio in FTD iPSCs treated with DCPIB was only partially rescued (mean 14.24 ± 0.65 s.e.m.) compared with control WT iPSCs (mean 22.06 ± 0.84), perhaps because of the short (24 h) treatment time. These findings underscore the association between increased pHlys and increased TDP-43 proteinopathy, suggesting targeting pHlys or using DCPIB as a potential therapeutic to reduce TDP-43 proteinopathy in FTD.

### FTD iPSCs have increased nuclear TFEB but similar total TFEB protein abundance compared with WT iPSCs

TFEB, a member of the MiTF/TFE transcription factor family, regulates the transcription of coordinated lysosomal expression and regulation (CLEAR) genes involved in autophagy and lysosome biogenesis, including lysosomal membrane proteins and hydrolases (Cho and Hwang, 2012; Wang et al., 2018; Ojalvo-Pacheco et al., 2024). Under normal conditions, phosphorylated TFEB remains in the cytosol, but during starvation or lysosomal stress it is dephosphorylated and translocated to the nucleus, activating lysosomal biogenesis (Cho and Hwang, 2012; Shariq et al., 2024). Immunolabeling for TFEB showed a greater nuclear to cytoplasmic ratio in FTD compared with WT iPSCs (Figure 5A–C), which is consistent with our findings that FTD iPSCs had more larger lysosomes compared with WT (Figure 1F) and lysosomal enlargement activating TFEB (Wang et al., 2018). Cytosolic TFEB in FTD iPSCs, however, was not notably in aggregates (Figure 5B) like we observed with TDP-43 (Figure 4B). Immunoblotting showed that FTD iPSCs had less phosphorylated TFEB than WT, but the total TFEB levels were similar (Figure 5D), consistent with phosphorylated TFEB being retained in the cytoplasm. Our findings are in agreement with previous reports indicating that loss of C9orf72 function in HEK293T cells leads to nuclear translocation of TFEB (Ugolino et al., 2016; Wang et al., 2020). In contrast to our findings, however, decreased nuclear TFEB expression was reported in post-mortem brain samples of AD and ALS patients (Wang et al., 2016).

**FIGURE 5:**
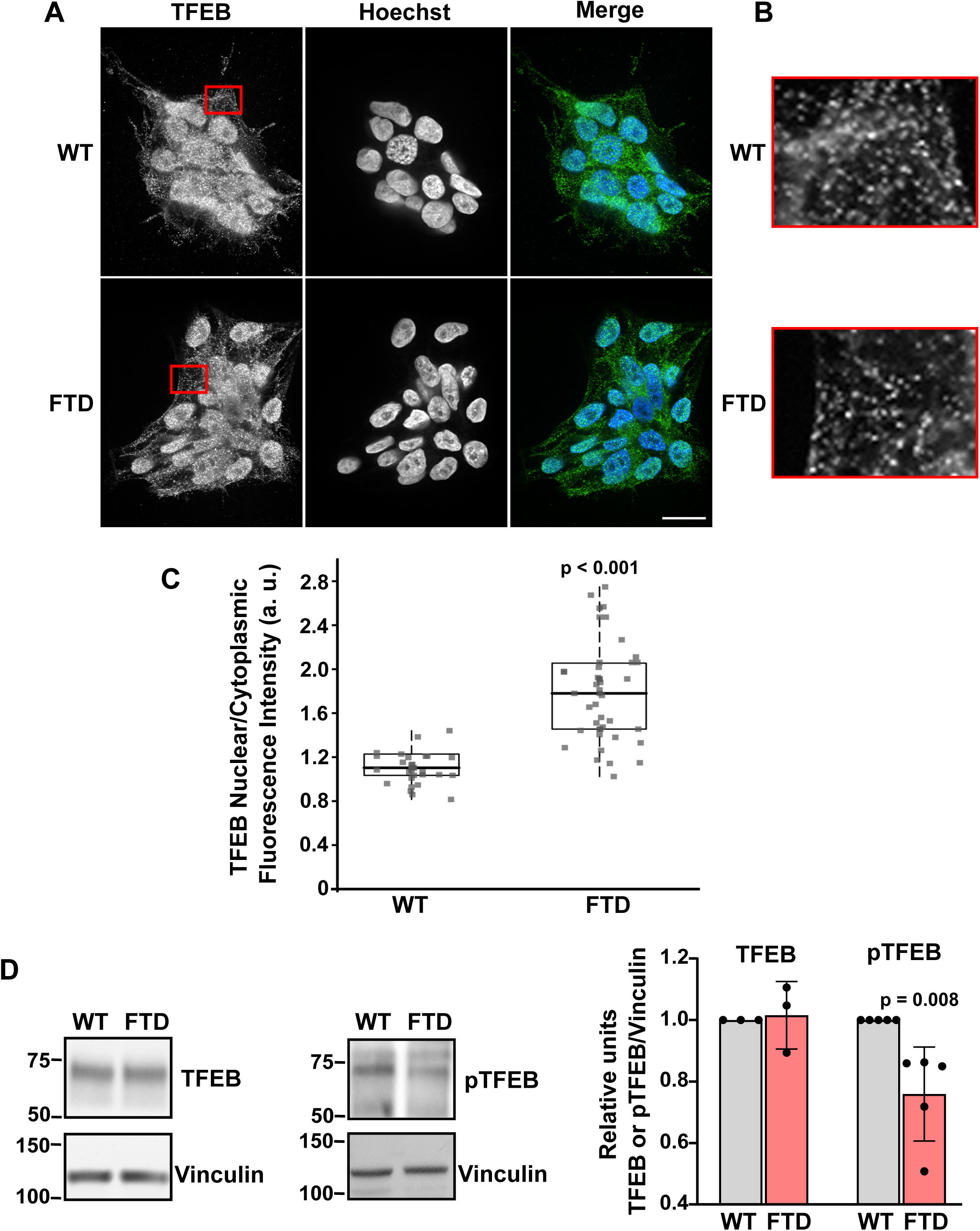
TFEB Localization and Expression. (A) Confocal images of TFEB immunolabeling in WT and FTD iPSCs and stained with Hoechst 33342. Scale bar 10 μm. (B) Higher magnification of the red-highlighted areas in (A) of TFEB immunolabeling. (C) TFEB nuclear/cytoplasmic ratios in WT and FTD iPSCs quantified from three separate cell preparations. Each dot represents the value from a single cell. (D) Quantification and representative immunoblot from three separate cell preparations for total TFEB and pTFEB with immunoblotting for vinculin as a loading control.

Lysosome biogenesis is also promoted by the activity of TFE3, a bHLH-leucine zipper transcription factor that, like TFEB, translocates from the cytoplasm to the nucleus when active and regulates autophagy (Jain et al., 2022; Mutvei et al., 2023; Ojalvo-Pacheco et al., 2024). However, unlike the increased nuclear TFEB we observed in FTD compared with WT iPSCs, we observed no difference in TFE3 localization between the two cell lines (Supplemental Figure S3A, B), determined by immunolabeling, or TFE3 abundance, determined by immunoblotting total cell lysates (Supplemental Figure S3C). The role of TFE3 in FTD is controversial. TFEB and TFE3 interact with different regulatory proteins and enzyme; therefore, alterations in TFEB-specific regulators, such as mTORC1 or calcineurin, may selectively impact TFEB activity without influencing TFE3. Additionally, mutations in genes involved in TFEB regulation, such as CLEAR network genes, may disrupt TFEB function without directly affecting TFE3. Despite both TFEB and TFE3 having a conserved transcriptional activation domain, TFEB aggregation is facilitated by the PrLD domain, which is absent in TFE3, and suggested as a target in Huntington’s disease (HD) (Ojalvo-Pacheco et al., 2024). Although C9orf72 mutations are associated with increased nuclear TFE3 (Wang et al., 2020), a general view is that dysregulated TFE3 localization may not be a hallmark of neurodegeneration (Ojalvo-Pacheco et al., 2024). Additionally, we saw no difference between FTD and WT iPSCs in the abundance of total or phosphorylated ULK1, a protein activated with autophagy stimulation (Amick and Ferguson, 2017; Root et al., 2021) (Supplemental Figure S3D). Our data indicate that in FTD iPSCs, TFEB regulation was distinct from changes in TFE3, ULK, and pULK, highlighting the complexity of signaling networks whereby C9orf72-G4C2 affects lysosome biology.

### Transcriptomic Profiling Reveals Dysregulated Pathways in FTD Compared with WT iPSCs

For unbiased insights on possible differences in gene expression and signaling networks in FTD compared with WT iPSCs we used RNA sequencing (RNA-seq). A Venn diagram revealed the number of shared and unique transcript levels between WT and FTDs (Figure 6A), with WT iPSCs having a total of 1,476 differentially expressed genes (DEGs) with an adjusted q-value of less than 0.05 after batch correction. Of these DEGs, 433 were unique to WT compared with FTD iPSCs. FTD iPSCs had 1,904 DEGs, with 861 unique DEGs. WT and FTD shared 1043 DEGs, which is almost 45% of their total number of DEGs (Figure 6A). A volcano plot, with each dot representing a gene, revealed increased and decreased transcripts in FTD compared with WT iPSCs (Figure 6B), which could potentially be used as biomarkers. DEGs (q-value < 0.05) are shown in blue, and we highlighted upregulated genes of interest in green and downregulated genes of interest in magenta based on literature meaning (Figure 6B). Compared with WT iPSCs, FTD iPSCs had increased expression of genes associated with synaptic and cytoskeletal processes, including moesin (MSN) (Beckmann et al., 2023) and presynaptic scaffolding RIM family members (RIMS1, RIMS2, RIMS3, RIMS4). Additionally, FTD IPSCs had increased expression of transcripts for synaptic regulation, such as CHGA and NNAT (Seredenina et al., 2011; Sathe et al., 2021; Podvin et al., 2022; Zou et al., 2022), as well as for inflammatory response, including MFGE8 (Almansoub et al., 2019) and IDO1 (Duan et al., 2020), which are linked to dementia. Similarly, FTD iPSCs had increased expression of several transcripts not previously associated with neurodegeneration, such as lysosome-associated H^+^-coupled amino acid transporters, SLC36A1 and SLC36A4 (Thwaites and Anderson, 2011; Hu et al., 2022), the lysosomal stress response protein LGALS3BP (Pan et al., 2014), and the neurotransmitter release transcripts belonging to SYT family (SYT4, SYT10 and SYT13) (Südhof, 2012).

**FIGURE 6:**
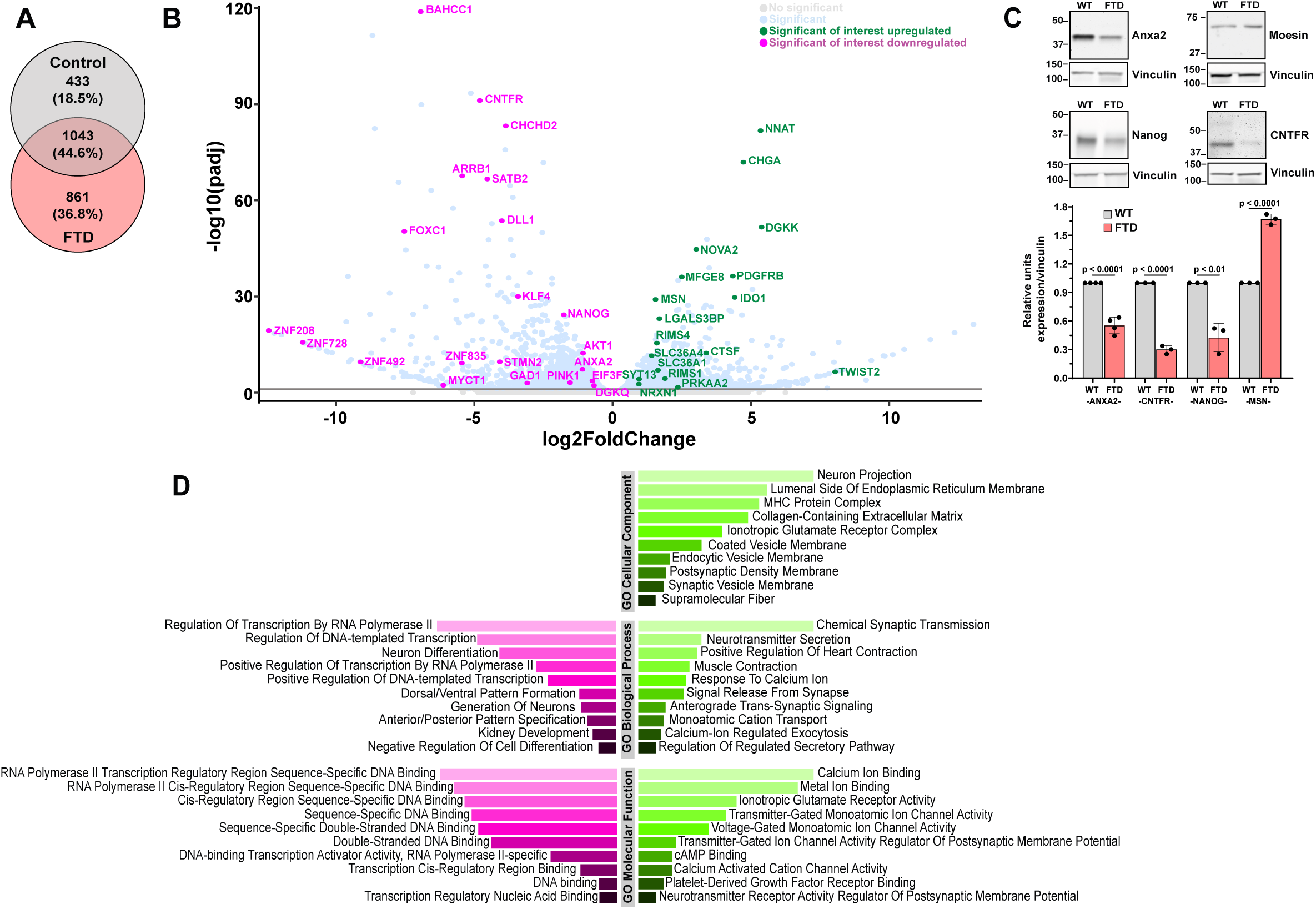
Transcriptomic Differences in WT and FTD iPSCs. (A) Venn diagram showing the number of shared and distinct DEGs in WT and FTD iPSCs. (B) Volcano plot showing the transcriptome fold-changes (beta values) in FTD compared with WT iPSCs. Each dot represents one gene, significantly changed genes (q-value < 0.05) indicated in blue. Among the significantly changed genes, those upregulated and of interest were highlighted in green while those downregulated and of interest were highlighted in magenta. (C) Immunoblots (above) and quantification (below) for the indicated proteins with vinculin used as a loading control. Quantification is presented as mean ± SEM of data from at least three separate cell preparations, with statistical significance determined by Student’s t-test. (D) Gene ontology molecular function, cellular component, and biological process enrichment analysis of upregulated DEGs uniquely indicated in FTD compared with WT iPSCs.

Gene transcripts decreased in FTD compared with WT iPSCs included some previously linked to ALS, FTD and/or PD, such as CHCHD2 (Imai et al., 2019), the ciliary neurotrophic factor receptor (CNTFR) (Deolankar et al., 2021), the neuronal development regulator SATB2 (Espuny-Camacho et al., 2013), four zinc-finger transcription factors (ZNF208, ZNF728, ZNF492, and ZNF835), as well as AKT1, which is related to lysosomal function and dysregulation (Paquette et al., 2021). Based on RNAseq findings, we confirmed by immunoblotting decreased expression in FTD compared with WT iPSCs of Annexin A2, a Ca^2+^ binding anti-apoptotic protein that regulates lysosome functions (Rentero et al., 2018; Quick et al., 2023) and CNTFR, which enhances neuronal survival (Yong et al., 2022) and has been associated with TDP-43 aggregation (Liao et al., 2022) (Fig. 6C). CNTFR is also downregulated in AD (Deolankar et al., 2021), although to our knowledge it has not been linked to FTD. In FTD iPSCs, we also confirmed, by immunoblotting, decreased expression of the pluripotency factor Nanog (Fig. 6C), which protects against neuronal toxicity (Chang et al., 2020). Immunoblotting confirmed that compared with WT iPSCs, FTD iPSCs have increased expression of the cytoskeletal linker MSN (Fig. 6C), which was recently shown to mediate tau-induced neurodegeneration (Beckmann et al., 2023). Moesin upregulation has also been reported in HIV-associated dementia in macaques, which mirrors the human disease (Gersten et al., 2009).

Moreover, we used gene ontology (GO) enrichment analysis of DEGs with the Enrichr database to reveal differences between FTD and WT iPSCs in cellular components (increases for neuron projections, glutamate receptor complex), biological processes (increases for neurotransmitter and synaptic transmission; decreases in neuron differentiation), and molecular functions (increases Ca^2+^ and neurotransmitter activity; decreases in DNA binding components (Figure 6D). Dysregulated Ca^2+^ homeostasis is associated with neurodegenerative conditions leading to neuronal loss (Hadi et al., 2024). In summary, these data revealed widespread transcriptomic differences in FTD compared with iPSCs, highlighting potential biomarkers as well as possible therapeutic targets.

## DISCUSSION

Motivated by the challenges and limitations of current FTD models, in this study we explored cellular pathologies using patient-derived iPSCs having a mutant C9ofr72-G4C2 expansion as an experimental model. We found that FTD iPSCs had disease hallmarks previously reported in neuronal models, including a disrupted lysosome biology of increased pHlys and decreased lysosomal hydrolase activity and specifically decreased cathepsin B activity, cytosolic protein aggregation, including TDP-43 proteinopathy, and increased nuclear TFEB. Furthermore, we found that pharmacologically decreasing pHlys attenuates TDP-43 proteinopathy in FTD iPSCs, highlighting the crucial role of lysosomal dysfunction in FTD. Moreover, RNAseq of FTD compared with WT iPSCs revealed dysregulated gene transcripts involved in calcium signaling, cell death, synaptic function, and neuronal development. Collectively, these findings underscore the potential of FTD iPSCs as a model for studying FTD cellular pathologies and for insights on signaling mechanisms regulated by C9orf72-G4C2.

We confirmed that pHlys was higher in FTD compared with WT iPSC by using two approaches, a genetically encoded pHlys biosensor pHLARE we developed (Webb et al., 2021) and the dye Oregon Green that was previously used to measure pHlys (Canton and Grinstein, 2017). A higher pHlys is also reported in FTD caused by progranulin haploinsufficiency (Tanaka et al., 2017; Elia et al., 2023), AD caused by presenilin 1 deficiency (Lee et al., 2010), and altered activity of the PD-risk protein TMEM175 (Hu et al., 2022). Collectively, these findings suggest that increased pHlys may be a hallmark of neurodegenerative disorders, as previously proposed (Nixon, 2017; Root et al., 2021). Further studies are needed to determine how C9orf72-G4C2 causes increased pHlys, with FTD iPSCs being a feasible model for resolving mechanisms. Although FTD and WT iPSCs had no difference in expression of TMEM175, which has been reported to regulate pHlys (Zhang et al., 2023), we cannot rule out possible differences in TMEM175 activity. We did find that DCPIB, a previously reported inhibitor of VRAC and swelling-activated chloride channels (Decher et al., 2001; Sardini et al., 2003), deceased pHlys with no effect on pHi. Treating FTD iPSCs with DCPIB also increased the reduced hydrolase activity and attenuated TDP-43 proteinopathy. Although not previously reported, of interest is whether DCPIB targets the lysosomal Cl-H exchanger ClC-7, which regulates pHlys (Graves et al., 2008; Coppola et al., 2023) and whether it might be considered as a therapeutic in treating FTD. Other possible regulators of increased pHlys in FTD cells are members of the SLC36 family of lysosomal proton-coupled amino acid transporters, as suggested by our RNAseq data indicating increased expression of SLC36A1 and SLC36A4. Alternatively, although receiving limited attention relative to neurodegenerative diseases, is whether a higher pHlys changes the expression or activity of SLC36 family members, and hence contributes to nutrient imbalances and dysregulated mTORC1 activity. Our finding that FTD iPSCs had a greater nuclear/cytoplasmic ratio of TFEB and less phosphorylated TFEB compared with WT iPSCs suggests decreased mTORC1 activity, although this remains to be confirmed.

Cytosolic accumulation and aggregates of TDP-43 are hallmarks of not only FTD (Greaves and Rohrer, 2019; Shao et al., 2022) but also other neurodegenerative disorders (Beckers and Van Damme, 2024) and lysosomal dysfunction may play a significant role in protein aggregate accumulation (Shao et al., 2022; Ojalvo-Pacheco et al., 2024). Of significance was our finding that TDP-43 proteinopathy was seen in FTD iPSCs in the absence of inducing neuronal differentiation. Reducing TDP-43 proteinopathy is a target for FTD therapeutics (Liao et al., 2022) and our results indicate that FTD iPSCs could be a tractable model toward achieving this goal and highlight the promise of targeting increased pHlys, which when reduced by DCPIB attenuated TDP-43 proteinopathy in FTD iPSCs.

Our RNAseq analysis highlights additional differences in FTD compared with WT iPSCs, particularly gene transcripts associated with synaptic regulation, including SYT family members, and neuronal development, including Nanog, SATB2 and ZNF transcription factors. RNAseq data confirmed dysregulated expression of several genes previously associated with FTD, ALS, or PD, including decreased expression of CNTFR, CHCHD2, AKT1, and NANOG {Deolankar, 2021 #918} (Deolankar et al., 2021; Imai et al., 2019; Paquette et al., 2021; Chang et al., 2020, (de Araujo et al., 2020)). Also previously linked to neurodegenerative disorders is increased expression of MSN (Beckmann et al., 2023). For RNAseq data, we confirmed dysregulated expression of CNTFR, ANXA2, Nanog and MSN by immunoblotting. Notably, in agreement with our immunoblotting data, RNAseq did not show differences in the expression of TMEM175, LAMP1, ULK, or Rab7.

Collectively, our findings support using iPSCs as a model for insights on the cell biology of FTD. Although C9orf72 binds to lysosomes (Amick and Ferguson, 2017), how wild-type and C9orf72-G4C2 regulate lysosome functions remains controversial (Root et al., 2021). The FTD iPSCs we used with variations to correct the C9orf72 mutation is an experimentally feasible model for resolving signaling networks and molecular mechanisms. Beyond C9orf72-G4C2 and FTD, iPSCs derived from patients having FTD and other neurodegeneration-causing mutations in MAPT/tau, progranulin, LRRK2 and PARK2 are available through the NINDS human cell repository (https://nindsgenetics.org/) and likely also useful for mechanistic insights on associated cell dysfunctions. Additionally, because of the time and cost-effective advantages of iPSCs compared with terminally differentiated neurons, we also suggest that iPSCs from affected patients are a feasible platform for drug screening to identify small molecules, for example that lower pHlys, with promise as therapeutics for treating FTD and other neurodegenerative disorders. Neurons derived from iPSCs carrying the C9orf72-G4C2 mutation have been used for screening small libraries (<2K bioactive compounds) (Czuppa et al., 2022). However, iPSCs offer greater feasibility for screening larger libraries, especially when combined with genetically encoded biosensors such as pHLARE, which enables longitudinal screening to identify small molecules that lower pHlys.

## MATERIALS AND METHODS

### Cell culture and transfection

Cell lines were obtained from the NINDS, including WT (NHCDR ID ND41866, Cell line ID NN0003921) and FTD (NHCDR ID ND50037, Cell line ID NN0000035) iPSCs. FTD iPSCs were generated from a disease-affected 48-year-old Caucasian male with a C9orf72-G4C2 expansion and WT iPSCs were generated from a non-affected 64-year-old, gender, and race matched individual. Cells were maintained in mTeSR Plus medium (STEMCELL Technologies) on Matrigel-coated dishes (Corning® Matrigel® hESC-Qualified Matrix) at 37°C and 5% CO_2_ and passaged by using Accutase to dissociate cells. After dissociation, cell suspensions were centrifuged at 1000 rpm for 5 min and the cell pellet was resuspended in mTeSR Plus medium supplemented with 10 μM of the Rho kinase inhibitor Y-27632 (MedChemExpress HY-10583) and plated on Matrigel-coated dishes. After 24 h, medium was replaced with mTeSR Plus lacking Y-27632. Mycoplasma contamination was tested 2×/year using medium obtained from cells maintained for 48 h by using a PCR Mycoplasma Detection Kit (abm #G238). Cell lines were authenticated commercially by IDEXX BioAnalytics.

### Stable expression of pHLARE

A mammalian expression plasmid with a CMV promoter encoding pHLARE was generated by our laboratory as previously reported (Webb et al., 2021), and is commercially available [Addgene #164477]. For stable expression in iPSCs, Gibson assembly was used to replace the CMV promoter with an EF-1α promoter. Confluent iPSCs, maintained as described above, were dissociated with Accutase, pelleted by centrifugation, and resuspended to 3-5×10^7^ cells/ml in MaxCyte EP buffer containing Y27632 (10 μM), transferring 50 µl of cell suspension into a separate microcentrifuge tube for each transfection to which 240 μg/ml of DNA/tube was added. For TALEN-mediated insertion to the AAVS1 locus we used a 4:3:3 ratio, donor construct:each of the site-specific TALENs (TALEN L [Addgene #59025] and TALEN R [Addgene #59026]). The cell suspension with DNA was transferred to R-50×8 Multi-well Processing Assembly electroporation cuvettes. Cells were electroporated using manufacture’s High Energy A549 protocol, and then allowed to recover for 30 min at 37 °C before plating cells Matrigel-coated dishes as described above. 72 h after electroporation, cells were sorted for mCherry fluorescence on a FACS Aria II flow cytometer (FACSDiva Software, BD Biosciences) at the UCSF Parnassus Flow Cytometry Core.

### Lysosome pH measurements

For live-cell imaging of pHlys, cells were plated on Matrigel-coated 35-mm MatTek dishes (MatTek Corporation; P35G-1.5-10-C) for 48 h. For imaging pHLARE, fluorescence ratios (Ex488 for sfGFP and Ex561 for mCherry) from cells in growth medium were acquired over 1 min, followed by acquiring fluorescence ratios after 5 min each in a nigericin-containing buffer (50 mM KPO_4_, 80 mM KCl, 1 mM MgCl) adjusted to 6.5 and 5.0 with HCl to calibrate ratios to pH values as previously described (Webb et al., 2021). Images were acquired using a customized spinning disk confocal (Yokogawa CSU-X1) on a Nikon Ti-E microscope with a 60Χ Plan TIRF 1.49 NA objective equipped with a Photometrics cMYO cooled CCD camera, as described (Stehbens et al., 2012). Lysosomal pH was also measured using ratiometric fluorescence imaging with the pH-sensitive dye Oregon Green™ 488 Dextran, 10,000 MW, Anionic, Lysine Fixable (Invitrogen, D7171), following previously described methods (Zhang et al., 2023) Twenty-four hours after plating, cells were washed, the medium was refreshed, and cells were incubated overnight with Oregon Green™ 488 at a concentration of 250 µg/mL. The following day, cells were washed three times and allowed a 3-hour dye-free chase in fresh medium before fixing and imaging. Fluorescence emissions were collected at 530 nm with excitation wavelengths of 440 nm and 488 nm, respectively. After each imaging measurement, in-situ pH calibration was performed by sequentially equilibrating cells with a series of pH-standard solutions for 5 minutes each, followed by imaging. The pH standards were prepared with 90 mM KCl, 25 mM MES, and 20 µM nigericin, adjusted to pH values of 4.1, 4.8, and 5.6. The ratio of fluorescence emission intensity of individual cells, acquired at excitation wavelengths of 488/440 nm after each pH treatment, was used to generate a standard curve, allowing the experimental fluorescence measurements to be interpolated into precise lysosomal pH values for each cell. Image acquisition settings that caused less than 5% photobleaching were used in all experiments. Images were analyzed using NIS Elements software and an analysis pipeline we developed as previously described (Webb et al., 2021). To determine lysosome size, we used the analysis pipeline program as previously described (Webb et al., 2021) that includes the area of lysosome objects. To evaluate the size of single lysosomes and not lysosome clusters, we used only the area of objects with a shape value between 0.9 and 1.0.

To experimentally change pHlys, 100 nM Bafilomycin A1 (Sigma-Aldrich; B1793) or 100 μM chloroquine (Sigma-Aldrich; C6628) was added to cells in growth medium for 2 h before pHLARE imaging, and 100 nM Torin-1 (Tocris Bioscience; 4247) or 10 μM DCPIB (Sigma-Aldrich; SML2692) was added to cells in growth medium for 24 h before pHLARE imaging as described above.

### Intracellular pH measurements (pHi)

Steady-state pHi was measured as previously described (Grillo-Hill et al., 2015; White et al., 2017). In brief, cells plated for 48 h in 24-well dishes were washed and incubated for 15 min in a NaHCO_3_ ^-^ containing buffer supplemented with 1 μM of the pH-sensitive dye BCECF. After washing in NaHCO_3_ ^-^ containing buffer, BCECF fluorescence ratios were measured using a SpectraMax fluorescence plate reader (Molecular Devices) and calibrated to pHi by treating cells at the end of each measurement with a potassium phosphate buffer containing the protonophore nigericin (10 μM) at pH 6.5 and 7.5, as previously described (Grillo-Hill et al., 2014). For measurements with DCPIB (10 μM), the inhibitor was added 24 h before pHi measurements and included in the BCECF-loading and pHi measurement buffers.

### Lysosomal hydrolytic activity assay

The overall lysosomal hydrolytic activity was measured using DQ™-Red BSA (Invitrogen, D12051). Cathepsin B activity was measured using the Magic Red Cathepsin B assay kit (ImmunoChemistry Technologies, #937). For both approaches, cells were plated on Matrigel-coated 35-mm MatTek dishes for 48 h and then incubated with DQ™-Red BSA (10 µg/ml) at 37 °C for 3 h or Magic Red (1:1000 dilution, following protocol instructions) at 37 °C for 30 minutes, followed by washing three times with PBS. Images were acquired (λ_exc_: 561 nm, λ_em_: 628 nm) using spinning disk confocal microscopy as described above. NIS Elements software (Nikon) was used to analyze the images, which included fluorescence intensity with subtracted background fluorescence. For intensity measurements, fluorescence intensity was quantified from multiple regions of interest (ROIs) in the Z section with the highest intensity for each junction, using a maximum intensity projection. Data were normalized within each biological replicate to the mean intensity for all control values. Statistical analyses were performed using Prism (GraphPad Software).

### PROTEOSTAT® Aggresome Detection Reagent

Protein aggregation was measured in cells plated for 48 h on Matrigel-coated 35-mm MatTek dishes. Cells were fixed in PBS containing 4% paraformaldehyde for 10 min at RT, and after washing were stained with 1/2000X diluted ProteoStat Aggresome Detection Kit (Enzo Life Sciences Inc.) according to the manufacturer’s protocol and with Hoechst 33342 (1 mg/ml) (Dojindo) to label nuclei for 30 min at room temperature in the dark. After washing with PBS, images were acquired with a Nikon Ti-E microscope with a 60Χ Plan TIRF 1.49 NA objective equipped with a Photometrics cMYO cooled CCD camera and analyzed using ImageJ software. To experimentally change protein aggregation, 100 nM Bafilomycin A1 (Sigma-Aldrich; B1793) was added to cells in growth medium for 2 h before fixing.

### Immunolabeling

Cells plated on Matrigel-coated 35-mm MatTek dishes for 48 h were washed 1X in PBS and fixed in PBS containing 4% paraformaldehyde for 10 min at RT. After washing in PBS, cells were permeabilized with 0.1% TritonX-100 in PBS for 10 min at RT, washed, and then incubated for 60 min in block buffer of PBS containing 5% FBS. Cells were then incubated with primary antibodies (Table 1) at 1:200 in PBS containing 0.01% Triton-X100 overnight at 4°C. After washing 3X with PBS, 10-min each wash, cells were incubated with fluorescent-conjugated secondary antibodies (1:200) (Jackson ImmunoResearch Laboratories) in PBS for 60 min at RT, and then washed 3X with PBS, 10-min each wash. For Hoechst 33342 staining, dye (1:10,000) was added to the second PBS wash. Fluorescence images were acquired by spinning disk confocal microscopy as described for Proteostat staining and analyzed using NIS Elements software. For analysis of nuclear/cytoplasmic ratios, for each cell analyzed, mean fluorescence intensities were obtained from 3 drawn ROI in the cytoplasm and 3 ROI in the nucleus, the latter indicated by Hoechst staining.

**Table 1.** Antibodies used.

| Antibodies | Source | Catalog # |
| --- | --- | --- |
| Actin C4 | Millipore | ZMS10004 |
| Annexin A2 | Abcam | ab41803 |
| Cathepsin B | Abcam | EPR4323 |
| CNTFR | Proteintech | 10796-1-AP |
| E-cadherin (HECD-1) | Takara | M106 |
| Lamp1 | Cell Signaling | 9091S |
| Moesin | Cell Signaling | 3105 |
| Nanog | Abcam | EPR2027(2) |
| OCT4 | Santa Cruz Biotech | SC-5279 |
| Rab7 | Cell Signaling | 9367T |
| SOX2 | Cell Signaling | 3579T |
| TDP-43 | Proteintech | 80002-1-RR |
| TFE3 | Proteintech | 14480-1-AP |
| TFEB | Invitrogen | PA1-31552 |
| pTFEB | Cell Signaling | 37681S |
| TMEM175 | Proteintech | 19925-1-AP |
| ULK1 | Cell Signaling | 8054S |
| pULK1 | Cell Signaling | 14202S |
| Vinculin | Millipore | V9131 |

### Immunoblotting

Total lysates were obtained from cells plated on Matrigel-coated 6 WP for 48 h. Cells were washed 3X in cold PBS on ice, lysed in 100 ul/well RIPA buffer (50 mM Tris-HCL, 150 mM NaCl, 1 mM EDTA, 1% NP-40, 0.5% deoxycholate, 0.1% SDS supplemented with protease inhibitors PEFA) while rotating at 4 °C for 10 min, scrapped into microfuge tubes, and a post-nuclear supernatant was collected after centrifugation at 12,000 rpm for 5 min at 4 °C. Proteins in lysates were separated by SDS–PAGE and transferred to polyvinylidene difluoride membranes. The membranes were incubated in blocking buffer (TBS containing 0.1% Tween [TBSTw] containing 5% BSA or 5% non-fat milk for 1 h and then incubated with primary antibodies (Table 1) (1:1000 dilution in TBSTw) overnight at 4 °C. After washing, membranes were incubated with peroxidase–conjugated secondary antibodies (Jackson ImmunoResearch Laboratories) in TBSTw for 1h at RT, and bound antibodies were developed by enhanced chemiluminescence using SuperSignal West Femto (Thermo Fisher Scientific) and imaged using an Alpha Innotech FluorChem Q (Alpha Innotech). Images were analyzed using ImageJ software.

### RNA sequencing

RNA was extracted from iPSCs plated for 48 h on a Matrigel-coated 6 WP with the RNAeasy Micro Kit (Qiagen #74004) according to the manufacturer’s protocol with the following modifications: after disrupting the cells with RLT buffer, cell suspension was transferred into a QIAshredder spin column (Qiagen #79656) and centrifuge. The lysate was transferred to a QIAshredder spin column and centrifuged at 13,000 rpm for 2 min. The RNA concentration and purity were measured using SpectraMax QuickDrop UV-Vis Spectrophotometer (Molecular Devices, #7546). Libraries were prepared with the RNA Library Prep Kit (Illumina) and paired-end sequencing was completed using the NovaSeq 6000 (Illumina) with 3 samples/lane. The raw RNA-Seq reads were analyzed by Novogene (Beijing, China). Functional enrichment analysis of differentially expressed genes was performed using Gene Ontology (GO) and Kyoto Encyclopedia of Genes and Genomes (KEGG) pathway analysis. The results were visualized using R packages such as ggplot2.

## Abbreviations

AD: Alzheimer’s Disease
ALS: Amyotrophic Lateral Sclerosis
ANXA2: annexin 2
CNTFR: Ciliary neurotrophic factor receptor
DCPIB: 4-(2-(6,7-Dichloro-2-cyclopentylindan-1-on-5-yl)ethyl)benzene-1,3-diol
FTD: Frontotemporal Dementia
GRN: progranulin
iPSCs: induced pluripotent stem cells
pHi: intracellular pH
LAMP1: Lysosomal Associated Membrane Protein 1
MSN: moesin
mTORC1: mammalian target of rapamycin complex 1
NINDS: National Institute of Neurological Disorders and Stroke
pHi: intracellular (cytosolic) pH
pHlys: lysosomal pH
PD: Parkinson’s disease
TDP-43: TAR DNA-binding protein 43
TMEM175: Transmembrane protein 175

## Data and code availability

RNA-seq data generated during this study have been deposited in Gene Expression Omnibus (https://www.ncbi.nlm.nih.gov/geo/). The accession number for the data reported in this paper is GEO: GSE273025. The overall project RNA-seq raw data is available for public download in the NCBI database under accession code PRJNA1121636. Software/packages used to analyze the dataset were freely available.

## Data presentation and statistical analysis

Sample sizes and p values were reported in the corresponding figure legends. Box-and-whisker plots were generated using Analyse-It for Microsoft Excel and show median, first, and third quartile, observations within 1.5 times the interquartile range, and all individual data points. Scatter dot plots and statistical analyses were generated in GraphPad Prism 10. Data distribution was tested to be normal. Unless otherwise noted, graphs show mean ± SEM. Statistical significance was determined using either statistical analysis by Tukey–Kramer HSD, two-way ANOVA, or unpaired t-test. Images were representative of at least three independent experiments. Experiments were not randomized, and the investigators were not blinded during data analysis.

## AUTHOR CONTRIBUTIONS

D.L.B. conceived the hypothesis. D.L.B. and S.I.-T. designed the study, developed experiments, and obtained and analyzed data as well as assembled figures and wrote the manuscript.

## ACKNOWLEDGMENTS

This work was supported by awards from the Alliance for Therapeutics in Neuroscience (ATN) Genentech (no. 4301), the UCSF Catalyst Program, and the National Institutes of Health (NIH CA197855).

## REFERENCES

Ahmed, M., Spicer, C., Harley, J., Taylor, J. P., Hanna, M., Patani, R. and Greensmith, L. (2023). Amplifying the Heat Shock Response Ameliorates ALS and FTD Pathology in Mouse and Human Models. Mol Neurobiol 60, 6896–6915.

Almansoub, H., Tang, H., Wu, Y., Wang, D. Q., Mahaman, Y. A. R., Wei, N., Liu, D. et al. (2019). Tau Abnormalities and the Potential Therapy in Alzheimer’s Disease. J Alzheimers Dis 67, 13–33.

Amick, J. and Ferguson, S. M. (2017). C9orf72: At the intersection of lysosome cell biology and neurodegenerative disease. Traffic 18, 267–276.

Babic Leko, M., Zupunski, V., Kirincich, J., Smilovic, D., Hortobagyi, T., Hof, P. R. and Simic, G. (2019). Molecular Mechanisms of Neurodegeneration Related to C9orf72 Hexanucleotide Repeat Expansion. Behav Neurol 2019, 2909168.

Bandyopadhyay, D., Cyphersmith, A., Zapata, J. A., Kim, Y. J. and Payne, C. K. (2014). Lysosome transport as a function of lysosome diameter. PLoS One 9, e86847.

Beckers, J., Tharkeshwar, A. K., Fumagalli, L., Contardo, M., Van Schoor, E., Fazal, R., Van Damme, P. et al. (2023). A toxic gain-of-function mechanism in C9orf72 ALS impairs the autophagy-lysosome pathway in neurons. Acta Neuropathol Commun 11, 151.

Beckers, J. and Van Damme, P. (2024). Toxic gain-of-function mechanisms in C9orf72 ALS-FTD neurons drive autophagy and lysosome dysfunction. Autophagy 1–3.

Beckmann, A., Ramirez, P., Gamez, M., Gonzalez, E., De Mange, J., Bieniek, K. F., Frost, B. et al. (2023). Moesin is an effector of tau-induced actin overstabilization, cell cycle activation, and neurotoxicity in Alzheimer’s disease. iScience 26, 106152.

Canton, J. and Grinstein, S. (2017). Measuring Phagosomal pH by Fluorescence Microscopy. Phagocytosis and Phagosomes - Methods and Protocols. R. Botelho. Department of Chemistry and Biology - Ryerson University -Toronto, ON, Canada, Springer.

Casey, J. R., Grinstein, S. and Orlowski, J. (2010). Sensors and regulators of intracellular pH. Nat. Rev. Mol. Cell Biol. 11, 50–61.

Chang, C. C., Li, H. H., Tsou, S. H., Hung, H. C., Liu, G. Y., Korolenko, T. A., Lin, C. L. et al. (2020). The Pluripotency Factor Nanog Protects against Neuronal Amyloid beta-Induced Toxicity and Oxidative Stress through Insulin Sensitivity Restoration. Cells 9,

Chew, J., Gendron, T. F., Prudencio, M., Sasaguri, H., Zhang, Y. J., Castanedes-Casey, M., Petrucelli, L. et al. (2015). Neurodegeneration. C9ORF72 repeat expansions in mice cause TDP-43 pathology, neuronal loss, and behavioral deficits. Science 348, 1151–1154.

Chin, M. Y., Ang, K. H., Davies, J., Alquezar, C., Garda, V. G., Rooney, B., Kao, A. W. et al. (2022). Phenotypic Screening Using High-Content Imaging to Identify Lysosomal pH Modulators in a Neuronal Cell Model. ACS Chem Neurosci 13, 1505–1516.

Cho, S. and Hwang, E. S. (2012). Status of mTOR activity may phenotypically differentiate senescence and quiescence. Mol Cells 33, 597–604.

Colacurcio, D. J. and Nixon, R. A. (2016). Disorders of lysosomal acidification-The emerging role of v-ATPase in aging and neurodegenerative disease. Ageing Res Rev 32, 75–88.

Coppola, M. A., Tettey-Matey, A., Imbrici, P., Gavazzo, P., Liantonio, A. and Pusch, M. (2023). Biophysical Aspects of Neurodegenerative and Neurodevelopmental Disorders Involving Endo-/Lysosomal CLC Cl(-)/H(+) Antiporters. Life (Basel) 13,

Czuppa, M., Dhingra, A., Zhou, Q., Schludi, C., König, L., Scharf, E., Edbauer, D. et al. (2022). Drug screen in iPSC-Neurons identifies nucleoside analogs as inhibitors of (G4C2)n expression in C9orf72 ALS/FTD. Cell Report 39, 110913.

de Araujo, M. E. G., Liebscher, G., Hess, M. W. and Huber, L. A. (2020). Lysosomal size matters. Traffic 21, 60–75.

Decher, N., Lang, H. J., Nilius, B., BruÈggemann, A., Busch, A. E. and Steinmeyer, K. (2001). DCPIB is a novel selective blocker of ICl,swell and prevents swelling-induced shortening of guinea-pig atrial action potential duration. Br J Pharmacol 134, 1467–1479.

DeJesus-Hernandez, M., Mackenzie, I. R., Boeve, B. F., Boxer, A. L., Baker, M., Rutherford, N. J., Rademakers, R. et al. (2011). Expanded GGGGCC hexanucleotide repeat in noncoding region of C9ORF72 causes chromosome 9p-linked FTD and ALS. Neuron 72, 245–256.

Deolankar, S. C., Patil, A. H., Rex, D. A. B., Subba, P., Mahadevan, A. and Prasad, T. S. K. (2021). Mapping Post-Translational Modifications in Brain Regions in Alzheimer’s Disease Using Proteomics Data Mining. OMICS 25, 525–536.

Duan, Z., Zhang, S., Liang, H., Xing, Z., Guo, L., Shi, L., Yang, Q. et al. (2020). Amyloid beta neurotoxicity is IDO1-Kyn-AhR dependent and blocked by IDO1 inhibitor. Signal Transduct Target Ther 5, 96.

Elia, L., Herting, B., Alijagic, A., Buselli, C., Wong, L., Morrison, G., Finkbeiner, S. et al. (2023). Frontotemporal Dementia Patient Neurons With Progranulin Deficiency Display Protein Dyshomeostasis. bioRxiv

Esanov, R., Belle, K. C., van Blitterswijk, M., Belzil, V. V., Rademakers, R., Dickson, D. W., Zeier, Z. et al. (2016). C9orf72 promoter hypermethylation is reduced while hydroxymethylation is acquired during reprogramming of ALS patient cells. Exp Neurol 277, 171–177.

Espuny-Camacho, I., Michelsen, K. A., Gall, D., Linaro, D., Hasche, A., Bonnefont, J., Vanderhaeghen, P. et al. (2013). Pyramidal neurons derived from human pluripotent stem cells integrate efficiently into mouse brain circuits in vivo. Neuron 77, 440–456.

Fang, B., Wang, D., Huang, M., Yu, G. and Li, H. (2010). Hypothesis on the relationship between the change in intracellular pH and incidence of sporadic Alzheimer’s disease or vascular dementia. Int J Neurosci 120, 591–595.

Farg, M. A., Sundaramoorthy, V., Sultana, J. M., Yang, S., Atkinson, R. A., Levina, V., Atkin, J. D. et al. (2014). C9ORF72, implicated in amytrophic lateral sclerosis and frontotemporal dementia, regulates endosomal trafficking. Hum Mol Genet 23, 3579–3595.

Gallwitz, L., Bleibaum, F., Voss, M., Schweizer, M., Spengler, K., Winter, D., Saftig, P. et al. (2024). Cellular depletion of major cathepsin proteases reveals their concerted activities for lysosomal proteolysis. Cell Mol Life Sci 81, 227.

Gersten, M., Alirezaei, M., Marcondes, M. C., Flynn, C., Ravasi, T., Ideker, T. and Fox, H. S. (2009). An integrated systems analysis implicates EGR1 downregulation in simian immunodeficiency virus encephalitis-induced neural dysfunction (data accessible at NCBI GEO database GDS4214). Journal 29, 12467–12476.

Graves, A. R., Curran, P. K., Smith, C. L. and Mindell, J. A. (2008). The Cl-/H+ antiporter ClC-7 is the primary chloride permeation pathway in lysosomes. Nature 453, 788–792.

Greaves, C. V. and Rohrer, J. D. (2019). An update on genetic frontotemporal dementia. J Neurol. 266, 2075–2086.

Grillo-Hill, B. K., Choi, C., Jimenez-Vidal, M. and Barber, D. L. (2015). Increased H^+^ efflux is sufficient to induce dysplasia and necessary for viability with oncogene expression. Elife 4,

Hadi, F., Mortaja, M. and Hadi, Z. (2024). Calcium (Ca^2+^) hemostasis, mitochondria, autophagy, and mitophagy contribute to Alzheimer’s disease as early moderators. Cell Biochem Funct 42, e4085.

Halcrow, P. W., Geiger, J. D. and Chen, X. (2021). Overcoming Chemoresistance: Altering pH of Cellular Compartments by Chloroquine and Hydroxychloroquine. Front Cell Dev Biol 9, 627639.

Halliday, G. M., Holton, J. L., Revesz, T. and Dickson, D. W. (2011). Neuropathology underlying clinical variability in patients with synucleinopathies. Acta Neuropathol 122, 187–204.

Hu, M., Li, P., Wang, C., Feng, X., Geng, Q., Chen, W., Xu, H. et al. (2022). Parkinson’s disease-risk protein TMEM175 is a proton-activated proton channel in lysosomes. Cell 185, 2292–2308 e2220.

Hu, M., Zhou, N., Cai, W. and Xu, H. (2022). Lysosomal solute and water transport. J Cell Biol 221,

Imai, Y., Meng, H., Shiba-Fukushima, K. and Hattori, N. (2019). Twin CHCH Proteins, CHCHD2, and CHCHD10: Key Molecules of Parkinson’s Disease, Amyotrophic Lateral Sclerosis, and Frontotemporal Dementia. Int J Mol Sci 20,

Irwin, D. J., Cairns, N. J., Grossman, M., McMillan, C. T., Lee, E. B., Van Deerlin, V. M., Trojanowski, J. Q. et al. (2015). Frontotemporal lobar degeneration: defining phenotypic diversity through personalized medicine. Acta Neuropathol 129, 469–491.

Jain, V., Bose, S., Arya, A. K. and Arif, T. (2022). Lysosomes in Stem Cell Quiescence: A Potential Therapeutic Target in Acute Myeloid Leukemia. Cancers (Basel) 14,

Kos, J., Mitrovic, A., Perisic Nanut, M. and Pislar, A. (2022). Lysosomal peptidases-intriguing roles in cancer progression and neurodegeneration. FEBS Open Bio 12, 708–738.

Lara Flores, E., Qi, A., Reilly, L., Santiana, M., Ward, M. and Cookson, M. (2023). iNDI Transcription Factor-NGN2 differentiation of human iPSC into cortical neurons Version 2. 83415.

Lee, J. H., Yu, W. H., Kumar, A., Lee, S., Mohan, P. S., Peterhoff, C. M., Nixon, R. A. et al. (2010). Lysosomal proteolysis and autophagy require presenilin 1 and are disrupted by Alzheimer-related PS1 mutations. Cell 141, 1146–1158.

Lee, S. and Huang, E. J. (2017). Modeling ALS and FTD with iPSC-derived neurons. Brain Res 1656, 88–97.

Levine, B. and Kroemer, G. (2008). Autophagy in the pathogenesis of disease. Cell 132, 27–42.

Liao, Y. Z., Ma, J. and Dou, J. Z. (2022). The Role of TDP-43 in Neurodegenerative Disease. Mol Neurobiol 59, 4223–4241.

Lines, G., Casey, J. M., Preza, E. and Wray, S. (2020). Modelling frontotemporal dementia using patient-derived induced pluripotent stem cells. Mol Cell Neurosci 109, 103553.

Ling, S. C., Polymenidou, M. and Cleveland, D. W. (2013). Converging mechanisms in ALS and FTD: disrupted RNA and protein homeostasis. Neuron 79, 416–438.

Lo, C. H. and Zeng, J. (2023). Defective lysosomal acidification: a new prognostic marker and therapeutic target for neurodegenerative diseases. Transl Neurodegener 12, 29.

Lopez-Herdoiza, M. B., Bauche, S., Wilmet, B., Le Duigou, C., Roussel, D., Frah, M., Latouche, M. et al. (2023). C9ORF72 knockdown triggers FTD-like symptoms and cell pathology in mice. Front Cell Neurosci 17, 1155929.

Majdi, A., Mahmoudi, J., Sadigh-Eteghad, S., Golzari, S. E., Sabermarouf, B. and Reyhani-Rad, S. (2016). Permissive role of cytosolic pH acidification in neurodegeneration: A closer look at its causes and consequences. J Neurosci Res 94, 879–887.

Marian, O. C., Teo, J. D., Lee, J. Y., Song, H., Kwok, J. B., Landin-Romero, R., Don, A. S. et al. (2023). Disrupted myelin lipid metabolism differentiates frontotemporal dementia caused by GRN and C9orf72 gene mutations. Acta Neuropathol Commun 11, 52.

Mizielinska, S., Grönke, S., Niccoli, T., Ridler, C. E., Clayton, E. L., Devoy, A., Isaacs, A. M. et al. (2014). C9orf72 repeat expansions cause neurodegeneration in Drosophila through arginine-rich proteins. Science 345, 1192–1194.

Mutvei, A. P., Nagiec, M. J. and Blenis, J. (2023). Balancing lysosome abundance in health and disease. Nat Cell Biol 25, 1254–1264.

Neumann, M. and Mackenzie, I. R. A. (2019). Review: Neuropathology of non-tau frontotemporal lobar degeneration. Neuropathol Appl Neurobiol 45, 19–40.

Neumann, M., Sampathu, D. M., Kwong, L. K., Truax, A. C., Micsenyi, M. C., Chou, T. T., M.-Y., L. V. et al. (2006). Ubiquitinated TDP-43 in Frontotemporal Lobar Degeneration and Amyotrophic Lateral Sclerosis. Science 314, 130–133.

Nixon, R. A. (2017). Amyloid precursor protein and endosomal-lysosomal dysfunction in Alzheimer’s disease: inseparable partners in a multifactorial disease. FASEB J 31, 2729–2743.

Nixon, R. A. (2020). The aging lysosome: An essential catalyst for late-onset neurodegenerative diseases. Biochim Biophys Acta Proteins Proteom 1868, 140443.

Ojalvo-Pacheco, J., Yakhine-Diop, S. M. S., Fuentes, J. M., Paredes-Barquero, M. and Niso-Santano, M. (2024). Role of TFEB in Huntington’s Disease. Biology 13,

Pan, S., Chen, R., Tamura, Y., Crispin, D. A., Lai, L. A., May, D. H., Brentnall, T. A. et al. (2014). Quantitative glycoproteomics analysis reveals changes in N-glycosylation level associated with pancreatic ductal adenocarcinoma. J Proteome Res 13, 1293–1306.

Paquette, M., El-Houjeiri, L., L, C. Z., Puustinen, P., Blanchette, P., Jeong, H., Pause, A. et al. (2021). AMPK-dependent phosphorylation is required for transcriptional activation of TFEB and TFE3. Autophagy 17, 3957–3975.

Podvin, S., Jiang, Z., Boyarko, B., Rossitto, L. A., O’Donoghue, A., Rissman, R. A. and Hook, V. (2022). Dysregulation of Neuropeptide and Tau Peptide Signatures in Human Alzheimer’s Disease Brain. ACS Chem Neurosci 13, 1992–2005.

Qi, Y., Zhang, X. J., Renier, N., Wu, Z., Atkin, T., Sun, Z., Studer, L. et al. (2017). Combined small-molecule inhibition accelerates the derivation of functional cortical neurons from human pluripotent stem cells. Nat Biotechnol 35, 154–163.

Quick, J. D., Silva, C., Wong, J. H., Lim, K. L., Reynolds, R., Barron, A. M., Lo, C. H. et al. (2023). Lysosomal acidification dysfunction in microglia: an emerging pathogenic mechanism of neuroinflammation and neurodegeneration. J Neuroinflammation 20, 185.

Rademakers, R., Neumann, M. and Mackenzie, I. R. (2012). Advances in understanding the molecular basis of frontotemporal dementia. Nat Rev Neurol 8, 423–434.

Raitano, S., Ordovas, L., De Muynck, L., Guo, W., Espuny-Camacho, I., Geraerts, M., Verfaillie, C. M. et al. (2015). Restoration of progranulin expression rescues cortical neuron generation in an induced pluripotent stem cell model of frontotemporal dementia. Stem Cell Reports 4, 16–24.

Ratti, A., Gumina, V., Lenzi, P., Bossolasco, P., Fulceri, F., Volpe, C., Colombrita, C. et al. (2020). Chronic stress induces formation of stress granules and pathological TDP-43 aggregates in human ALS fibroblasts and iPSC-motoneurons. Neurobiol Dis 145, 105051.

Rentero, C., Blanco-Munoz, P., Meneses-Salas, E., Grewal, T. and Enrich, C. (2018). Annexins-Coordinators of Cholesterol Homeostasis in Endocytic Pathways. Int J Mol Sci 19,

Renton, A. E., Majounie, E., Waite, A., Simon-Sanchez, J., Rollinson, S., Gibbs, J. R., Traynor, B. J. et al. (2011). A hexanucleotide repeat expansion in C9ORF72 is the cause of chromosome 9p21-linked ALS-FTD. Neuron 72, 257–268.

Root, J., Merino, P., Nuckols, A., Johnson, M. and Kukar, T. (2021). Lysosome dysfunction as a cause of neurodegenerative diseases: Lessons from frontotemporal dementia and amyotrophic lateral sclerosis. Neurobiol Dis 154, 105360.

Sardini, A., Amey, J. S., Weylandt, K. H., Nobles, M., Valverde, M. A. and Higgins, C. F. (2003). Cell volume regulation and swelling-activated chloride channels. Biochim Biophys Acta 1618, 153–162.

Sathe, G., Albert, M., Darrow, J., Saito, A., Troncoso, J., Pandey, A. and Moghekar, A. (2021). Quantitative proteomic analysis of the frontal cortex in Alzheimer’s disease. J Neurochem 156, 988–1002.

Selvaraj, B. T., Livesey, M. R. and Chandran, S. (2017). Modeling the C9ORF72 repeat expansion mutation using human induced pluripotent stem cells. Brain Pathol 27, 518–524.

Seredenina, T., Gokce, O. and Luthi-Carter, R. (2011). Decreased striatal RGS2 expression is neuroprotective in Huntington’s disease (HD) and exemplifies a compensatory aspect of HD-induced gene regulation. PLoS One 6, e22231.

Shao, W., Todd, T. W., Wu, Y., Jones, C. Y., Tong, J., Jansen-West, K., Petrucelli, L. et al. (2022). Two FTD-ALS genes converge on the endosomal pathway to induce TDP-43 pathology and degeneration. Science 378, 94–99.

Shariq, M., Khan, M. F., Raj, R., Ahsan, N. and Kumar, P. (2024). PRKAA2, MTOR, and TFEB in the regulation of lysosomal damage response and autophagy. J Mol Med (Berl) 102, 287–311.

Shaw, M. P., Higginbottom, A., McGown, A., Castelli, L. M., James, E., Hautbergue, G. M., Ramesh, T. M. et al. (2018). Stable transgenic C9orf72 zebrafish model key aspects of the ALS/FTD phenotype and reveal novel pathological features. Acta Neuropathol Commun 6, 125.

Shi, Y., Lin, S., Staats, K. A., Li, Y., Chang, W. H., Hung, S. T., Ichida, J. K. et al. (2018). Haploinsufficiency leads to neurodegeneration in C9ORF72 ALS/FTD human induced motor neurons. Nat Med 24, 313–325.

Siddiqui, T. and Bhatt, L. K. (2023). Emerging autophagic endo-lysosomal targets in the management of Parkinson’s disease. Rev Neurol (Paris)

Smeele, P. H., Cesare, G. and Vaccari, T. (2024). ALS’ Perfect Storm: C9orf72-Associated Toxic Dipeptide Repeats as Potential Multipotent Disruptors of Protein Homeostasis. Cells 13,

Smith, J. R., Maguire, S., Davis, L. A., Alexander, M., Yang, F., Chandran, S., Pedersen, R. A. et al. (2008). Robust, persistent transgene expression in human embryonic stem cells is achieved with AAVS1-targeted integration. Stem Cells 26, 496–504.

Sonobe, Y., Lee, S., Krishnan, G., Gu, Y., Kwon, D. Y., Gao, F. B., Kratsios, P. et al. (2023). Translation of dipeptide repeat proteins in C9ORF72 ALS/FTD through unique and redundant AUG initiation codons. Elife 12,

Stehbens, S., Pemble, H., Murrow, L. and Wittmann, T. (2012). Imaging intracellular protein dynamics by spinning disk confocal microscopy. Methods Enzymol 504, 293–313.

Südhof, T. C. (2012). Calcium control of neurotransmitter release. Cold Spring Harb Perspect Biol 4, a011353.

Swift, I. J., Sjodin, S., Gobom, J., Brinkmalm, A., Blennow, K., Zetterberg, H., Sogorb-Esteve, A. et al. (2024). Differential patterns of lysosomal dysfunction are seen in the clinicopathological forms of primary progressive aphasia. J Neurol 271, 1277–1285.

Tanaka, Y., Ito, S. I., Honma, Y., Hasegawa, M., Kametani, F., Suzuki, G., Eto, M. et al. (2023). Dysregulation of the progranulin-driven autophagy-lysosomal pathway mediates secretion of the nuclear protein TDP-43. J Biol Chem 299, 105272.

Tanaka, Y., Suzuki, G., Matsuwaki, T., Hosokawa, M., Serrano, G., Beach, T. G., Nishihara, M. et al. (2017). Progranulin regulates lysosomal function and biogenesis through acidification of lysosomes. Hum Mol Genet 26, 969–988.

Tang, T., Jian, B. and Liu, Z. (2023). Transmembrane Protein 175, a Lysosomal Ion Channel Related to Parkinson’s Disease. Biomolecules 13,

Taylor, J. P., Hardy, J. and Fischbeck, K. H. (2002). Toxic Proteins in Neurodegenerative Disease. Science 296, 1991–1994.

Thoreen, C. C., Kang, S. A., Chang, J. W., Liu, Q., Zhang, J., Gao, Y., Gray, N. S. et al. (2009). An ATP-competitive mammalian target of rapamycin inhibitor reveals rapamycin-resistant functions of mTORC1. J Biol Chem 284, 8023–8032.

Thwaites, D. T. and Anderson, C. M. (2011). The SLC36 family of proton-coupled amino acid transporters and their potential role in drug transport. Br J Pharmacol 164, 1802–1816.

Todd, T. W., Shao, W., Zhang, Y. J. and Petrucelli, L. (2023). The endolysosomal pathway and ALS/FTD. Trends Neurosci 46, 1025–1041.

Tu, C., Ortega-Cava, C. F., Chen, G., Fernandes, N. D., Cavallo-Medved, D., Sloane, B. F., Band, H. et al. (2008). Lysosomal cathepsin B participates in the podosome-mediated extracellular matrix degradation and invasion via secreted lysosomes in v-Src fibroblasts. Cancer Res 68, 9147–9156.

Ugolino, J., Ji, Y. J., Conchina, K., Chu, J., Nirujogi, R. S., Pandey, A., Wang, J. et al. (2016). Loss of C9orf72 Enhances Autophagic Activity via Deregulated mTOR and TFEB Signaling. PLoS Genet 12, e1006443.

Valdez, C., Wong, Y. C., Schwake, M., Bu, G., Wszolek, Z. K. and Krainc, D. (2017). Progranulin-mediated deficiency of cathepsin D results in FTD and NCL-like phenotypes in neurons derived from FTD patients. Hum Mol Genet 26, 4861–4872.

Vo, Q. D., Saito, Y., Ida, T., Nakamura, K. and Yuasa, S. (2024). The use of artificial intelligence in induced pluripotent stem cell-based technology over 10-year period: A systematic scoping review. PLoS One 19, e0302537.

Wang, F., Gomez-Sintes, R. and Boya, P. (2018). Lysosomal membrane permeabilization and cell death. Traffic 19, 918–931.

Wang, H., Wang, R., Xu, S. and Lakshmana, M. K. (2016). Transcription Factor EB Is Selectively Reduced in the Nuclear Fractions of Alzheimer’s and Amyotrophic Lateral Sclerosis Brains. Neurosci J 2016, 4732837.

Wang, M., Wang, H., Tao, Z., Xia, Q., Hao, Z., Prehn, J. H. M., Ying, Z. et al. (2020). C9orf72 associates with inactive Rag GTPases and regulates mTORC1-mediated autophagosomal and lysosomal biogenesis. Aging Cell 19, e13126.

Webb, B. A., Aloisio, F. M., Charafeddine, R. A., Cook, J., Wittmann, T. and Barber, D. L. (2021). pHLARE: a new biosensor reveals decreased lysosome pH in cancer cells. Mol Biol Cell 32, 131–142.

White, A. J., Clark, K. A., Alexander, K. D., Ramalingam, N., Young-Pearse, T. L., Dettmer, U., Ho, G. P. H. et al. (2024). A stem cell-based assay platform demonstrates alpha-synuclein dependent synaptic dysfunction in patient-derived cortical neurons. NPJ Parkinsons Dis 10, 107.

White, K. A., Grillo-Hill, B. K. and Barber, D. L. (2017). Cancer cell behaviors mediated by dysregulated pH dynamics at a glance. J Cell Sci 130, 663–669.

Wolfe, D. M., Lee, J. H., Kumar, A., Lee, S., Orenstein, S. J. and Nixon, R. A. (2013). Autophagy failure in Alzheimer’s disease and the role of defective lysosomal acidification. Eur J Neurosci 37, 1949–1961.

Xu, G., Peng, H., Yao, R., Yang, Y. and Li, B. (2024). TFEB and TFE3 cooperate in regulating inorganic arsenic-induced autophagy-lysosome impairment and immuno-dysfunction in primary dendritic cells. Cell Biol Toxicol 40, 4.

Yong, J., Groeger, S., von Bremen, J. and Ruf, S. (2022). Ciliary Neurotrophic Factor (CNTF) and Its Receptors Signal Regulate Cementoblasts Apoptosis through a Mechanism of ERK1/2 and Caspases Signaling. Int J Mol Sci 23,

Zhang, J., Zeng, W., Han, Y., Lee, W. R., Liou, J. and Jiang, Y. (2023). Lysosomal LAMP proteins regulate lysosomal pH by direct inhibition of the TMEM175 channel. Mol Cell 83, 2524–2539 e2527.

Zou, H. Y., Guo, L., Zhang, B., Chen, S., Wu, X. R., Liu, X. D., Sun, S. et al. (2022). Aberrant miR-339-5p/neuronatin signaling causes prodromal neuronal calcium dyshomeostasis in mutant presenilin mice. J Clin Invest 132, e149160.

