## Supplementary Figures for "Patient-derived Induced Pluripotent Stem Cells as a Model to Study Frontotemporal Dementia Pathologies"

### Supplementary Figure 1

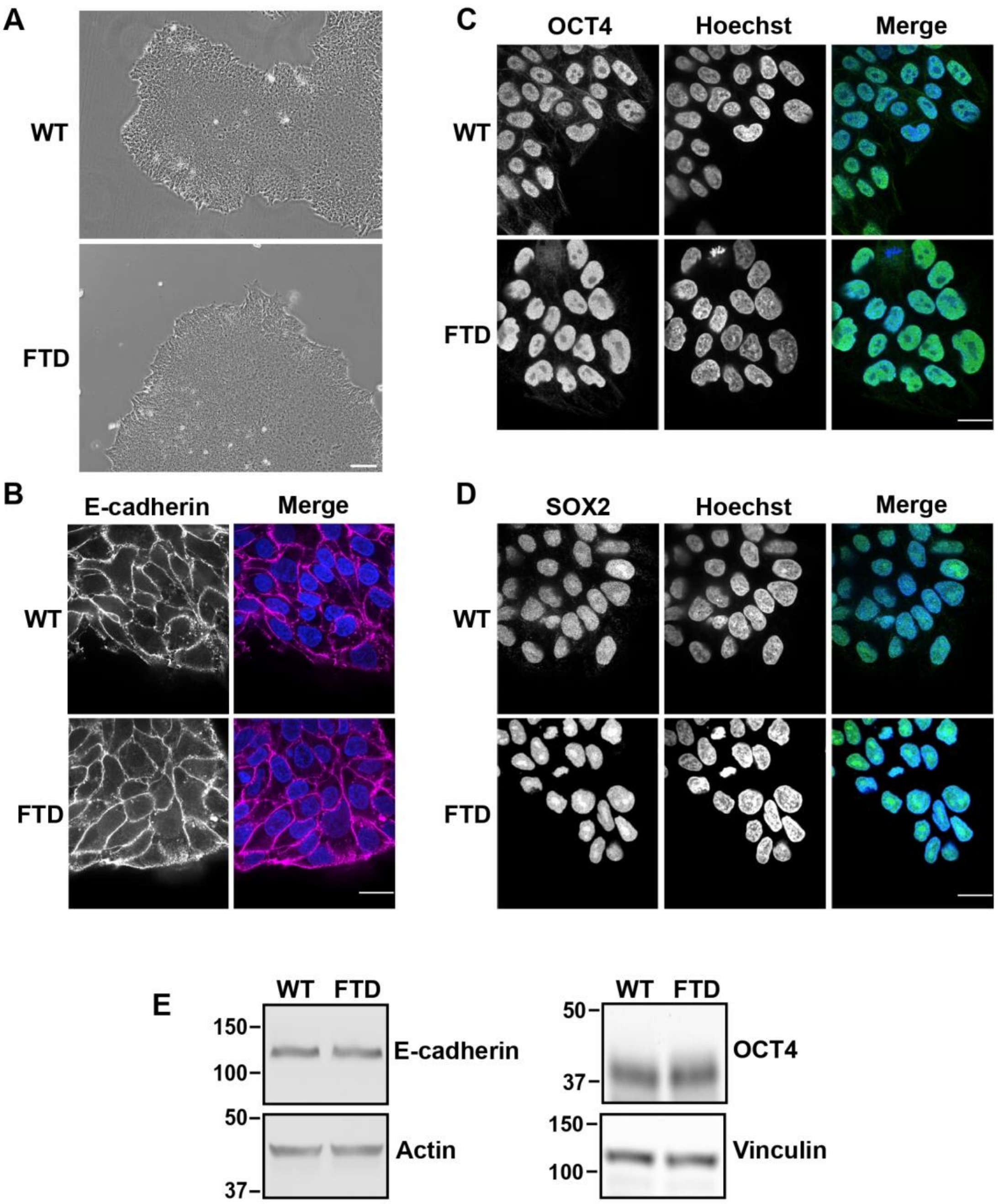


**FIGURE S1.** Characterization of iPSC lines. (A) Bright-field images of WT and FTD iPSC colonies. Scale bar 100 µm. Images were taken 48 h after removing Y-27632 (Rho-kinase inhibitor), allowing the formation of dense colonies. (B-D) Confocal images of immunolabeling for E-cadherin (B), OCT4 (C) and SOX2 (D) and Hoechst 33342 staining. Scale bar 10 µm. (E) Representative immunoblots for E-cadherin and OCT4 in WT and FTD iPSCs from three separate cell preparations.

#### Supplementary Figure 2

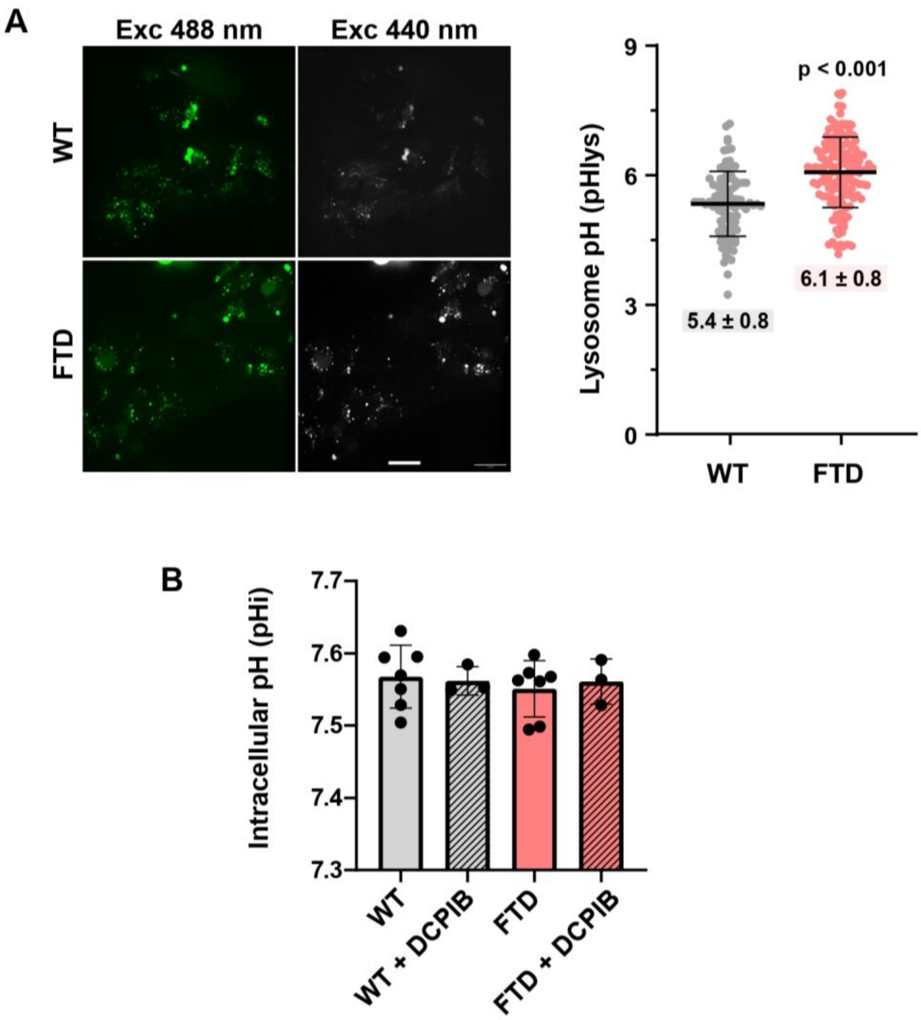


**FIGURE S2.** Lysosome and intracellular pH. (A) Confocal images of WT and FTD iPSCs to determine lysosome pH by using Oregon Green, showing excitation at 488 nm and 440 nm. Scale bar 20 μm. Quantified pHlys in WT and FTD iPSCs obtained from emission at 488/440 ratios calibrated with nigericin-containing buffers of known pH values. Each data point indicates an individual cell (WT 96 cells; FTD 150 cells) obtained from 3 separate cell preparations. Statistical analysis by Student's t-test comparison. (B) Intracellular (cytosolic) pH (pHi) determined in cells loaded with the pH-sensitive dye BCECF. Data show pHi for WT (grey) and FTD (red) iPSCs, with (striped bars) and without (solid bars) DCPIB treatment (10 μM, 18 h). Each dot represents the mean pHi value from three independent replicates performed on the same day. Error bars indicate standard deviation.

#### Supplementary Figure 3

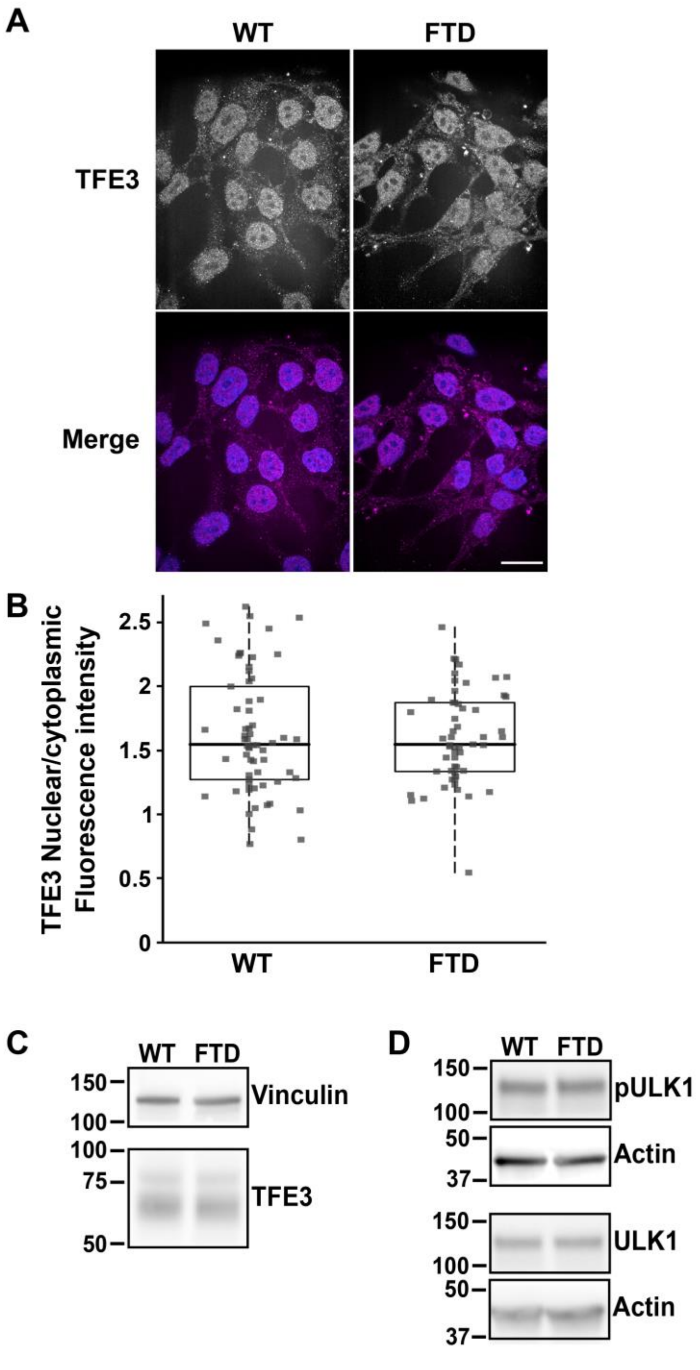


**FIGURE S3**. TFE3 and ULK expression. (A) Confocal images of TFE3 immunolabeling and merged with Hoechst 33342 staining in WT and FTD iPSCs. Scale bar 10 μm. (B) Quantification of nuclear/cytoplasmic ratios of TFE3 fluorescence intensities with data obtained from three separate cell preparations. (C, D) Representative immunoblots for TFE3 (C) and pULK1 and ULK1 (D) from three separate cell preparations.
